# Protein-protein interactions drive differences in the spatiotemporal dynamics of transcription factors NANOG and SOX2 in naïve pluripotent cells

**DOI:** 10.64898/2025.12.03.691924

**Authors:** Gökçe G. Agsu, Yuze Cao, Diyan Jain, Stanley E. Strawbridge, Sam Daly, Ziwei Zhang, Annabelle Wurmser, Kathryn Bowers, Igor Orsine De Almeida, Gergo Nikolenyi, Sebastian Berger, Mantas Jonaitis, Khalid Guma’a, Tekle Pauzaite, Smitha Maretvadakethope, Lawrence Bates, James A. Nathan, Brian D. Hendrich, Ruben Perez-Carrasco, Kevin J. Chalut, David Klenerman, Thorsten E. Boroviak, Steven F. Lee, David Holcman, Srinjan Basu

**Affiliations:** Department of Life Sciences, Imperial College London, London SW7 2AZ, UK; Cambridge Stem Cell Institute, University of Cambridge, Cambridge, CB2 0AW, UK; Group of Applied Mathematics and Computational Biology, Ecole Normale Supérieure, 75005 Paris, France; Department of Physiology, Development and Neuroscience, University of Cambridge, Cambridge, CB2 3DY, UK; Centre for Stem Cell Biology, School of Biosciences, University of Sheffield, Sheffield, S10 2TN, UK; Yusuf Hamied Department of Chemistry, University of Cambridge, Lensfield Road, Cambridge, CB2 1EW, UK; Cambridge Institute for Medical Research, University of Cambridge, Cambridge, CB2 0XY, UK; Department of Biochemistry, University of Cambridge, Cambridge, CB2 1GA, UK; Cambridge Institute of Therapeutic Immunology & Infectious Disease (CITIID), Department of Medicine, University of Cambridge, Cambridge, CB2 0AW, UK; MRC Human Genetics Unit, Institute of Genetics and Cancer, University of Edinburgh, Crewe Road South, Edinburgh EH4 2XU, UK; DAMPT, University of Cambridge, Cambridge, CB3 0XS, UK; Churchill College, Cambridge, CB30DS, UK

## Abstract

Maintenance of naïve pluripotency requires core transcription factors (TFs) like SOX2 and auxiliary TFs like NANOG, yet molecular mechanisms governing their intra-nuclear dynamics and DNA binding interactions remain unclear. Here, using high-density 3D single-molecule light-field microscopy combined with novel spatiotemporal analysis pipelines, we track SOX2 and NANOG dynamics in live cells. Despite lower protein abundance, NANOG displays a similar chromatin-bound fraction to SOX2. This arises partially because, while both TFs undergo frequent transient non-specific binding interactions (∼0.5-0.7s), NANOG exhibits more stable specific binding (∼25s vs ∼16s). Both TFs also assemble into phase-separated domains of ∼400 nm containing both freely diffusing and chromatin-bound proteins, which further influences their dynamics. Strikingly, NANOG’s protein-protein interaction domain markedly increases chromatin residence time (>5-fold) and the size of these phase-separated domains. Our work uncovers how NANOG and SOX2 stabilise gene regulatory networks that maintain naïve pluripotency while providing quantitative pipelines for dissecting spatiotemporal TF dynamics.

## Introduction

Naïve pluripotent cells possess the unique ability to differentiate into all the three germ layers that make up the mammalian body and therefore hold huge potential for regenerative medicine. This naïve pluripotent state is commonly investigated using mouse embryonic stem cells (ESCs) derived from the inner cell mass of pre-implantation blastocysts and cultured under defined self-renewal conditions that mimic naïve pluripotent cells *in vivo* (Ying et al. 2008; Evans and Kaufman 1981; Martin 1981). ESCs have been used to identify transcription factors (TFs) that define the gene regulatory network of naïve pluripotent cells (Chen et al. 2008; Dunn et al. 2014; Niwa 2018). In this network, OCT4 and SOX2 are core irreplaceable TFs: their depletion causes ESCs to exit pluripotency and undergo differentiation (Chen et al. 2008; Dunn et al. 2014; Niwa 2018). Meanwhile NANOG, ESRRB and KLF4 act as auxiliary TFs: although not required for ESC self-renewal, they stabilise pluripotency when overexpressed and increase the propensity for differentiation when depleted (Chambers et al. 2007; Silva et al. 2009; Mitsui et al. 2003; Chambers et al. 2003).

The activity of these core and auxiliary TFs depends not only on their expression levels but also on how they move and interact with DNA across time and space within the nucleus. These spatiotemporal dynamics define how they control gene regulatory networks and therefore the stability of the pluripotent cell state. Indeed, subtle disruption to the chromatin binding kinetics or protein-protein interactions of these TFs cause cells to differentiate (Roberts et al. 2021; Gagliardi et al. 2013; Mullin et al. 2017). However, addressing the spatiotemporal dynamics of pluripotency TFs is challenging as they are intricately linked to their expression level and to their interactions with each other, RNA and chromatin within the nucleus – hard to recreate outside living cells (Mistri et al. 2022; Chen et al. 2014; Biddle, Nguyen, and Gunawardena 2019). For example, NANOG’s tryptophan-repeat (WR) domain forms interactions with SOX2, OCT4 and other NANOG molecules, interactions that are critical for stabilising naïve pluripotency (Choi et al. 2018; Mullin et al. 2008; Choi et al. 2022; Gagliardi et al. 2013; Mistri et al. 2022). The spatiotemporal dynamics of pluripotency TFs may also be influenced by membrane-less intra-nuclear compartments, or phase-separated domains (PSDs), including transcriptional condensates enriched for TFs, mediator and RNA polymerase (Wurmser and Basu 2022; Sabari et al. 2018; Boija et al. 2018). Indeed, there is evidence that these TFs can form condensates (Krainer et al. 2021; Wang et al. 2021; Choi et al. 2022; Boija et al. 2018), although studies have primarily been conducted using purified TFs *in vitro*, in fixed cells or in cells expressing TFs at exogenously high levels. It is therefore unclear if these condensates form within living cells at endogenous TF concentrations, especially since fixation itself alters condensate properties (Irgen-Gioro et al. 2022) and since the DNA binding sites of TFs also cluster in 3D, which can create high apparent local concentrations (Ma et al. 2018; Stevens et al. 2017; Liu et al. 2014).

3D single-molecule localisation microscopy (3D-SMLM) allows us to investigate the spatiotemporal dynamics of both the core and auxiliary pluripotency TFs in naïve pluripotent cells. Unlike approaches such as FRAP, it is ideal for probing low-expressing proteins such as the auxiliary TF NANOG. It also does not suffer from limitations of 2D-SMLM e.g. difficulties detecting freely diffusing TFs and underestimation of residence times for chromatin-bound TFs that bind for long periods. Moreover, it has previously been used successfully to reveal the chromatin binding kinetics of the core TFs OCT4 and SOX2 (Chen et al. 2014; Biddle, Nguyen, and Gunawardena 2019) although using culture conditions that represent a mixture of naïve and primed pluripotent cell states (Marks et al. 2012).

In this study, we implement a recently established 3D-SMLM approach called single-molecule light-field microscopy (SMLFM) (Sims et al. 2020; Daly et al. 2024) alongside novel trajectory analysis pipelines to compare the spatiotemporal dynamics of the core TF SOX2 and the auxiliary TF NANOG in naïve pluripotent cells. Our results reveal differences in expression level, chromatin binding kinetics and spatial clustering of SOX2 and NANOG that may account for the different roles played by these TFs. NANOG’s dynamics are influenced by its WR protein-protein interaction domain, which ensures robust binding of NANOG at the single-cell level despite its low expression levels while increasing the size of PSDs formed in cells.

## Results

### NANOG is expressed at lower levels than SOX2

To determine the spatiotemporal strategies adopted by NANOG and SOX2 in finding their binding sites, we first determined their relative expression levels. To do this, we generated heterozygous knock-in ESCs expressing NANOG-Halo and SOX2-Halo fusion proteins from their endogenous loci (**Fig.1A, Supp Fig.1A**). HaloTag proteins allowed stoichiometric labelling with the bright JFX_554_ fluorophore and this was used as a proxy to compare relative protein levels (Grimm et al. 2015; Los et al. 2008). This revealed that NANOG was expressed at ∼9-fold lower levels than SOX2 (p=0.04) (**Fig.1A-D**). To ensure that fusion proteins were functional and expressed at endogenous levels, we carried out RT-qPCR and quantitative immunofluorescence. This revealed that NANOG-Halo and SOX2-Halo protein expression were at endogenous levels, despite minor changes in RNA level (p=10^-4^ and p=0.015), and that proteins were functional with no changes in the RNA levels of NANOG/SOX2-dependent genes e.g. *Esrrb* (**Supp Fig.1B-E**). We therefore concluded that HaloTag has not affected the functionality of NANOG/SOX2 and that NANOG is expressed at 9x lower levels than SOX2.

**Figure 1:**
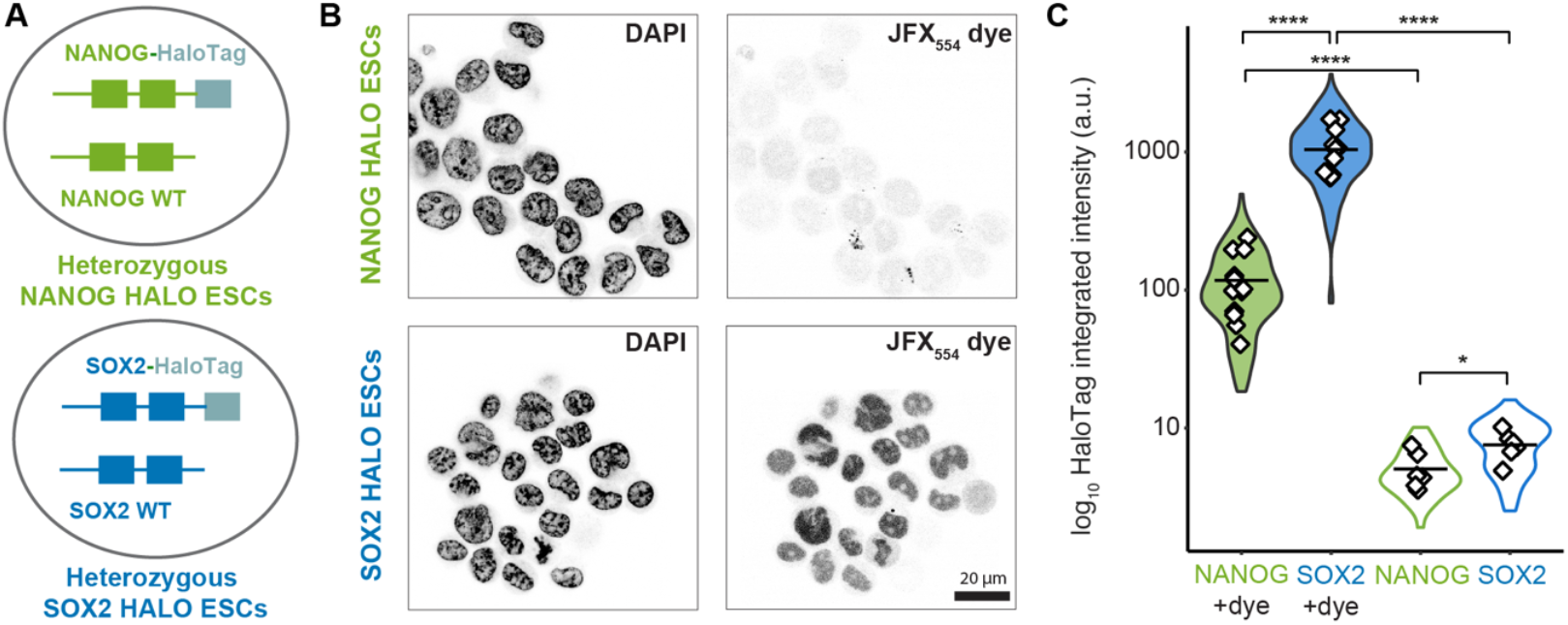
NANOG expression is ∼9 times lower than SOX2. **(A)** Schematic of heterozygous ESCs generated where one allele is labelled with HaloTag. **(B)** Representative images of DAPI and HaloTag-JFX_549_ intensity, a proxy for NANOG or SOX2 protein levels. **(C)** Violin plots showing log_10_ HaloTag integrated intensity distributions for NANOG-Halo and SOX2-Halo proteins with and without JFX_554_ dye (filled and hollow violin plots represent with and without dye respectively). [two-tailed unpaired t-test: NANOG+dye vs NANOG (no dye) (****p = 10^-5^), SOX2+dye vs SOX2 (no dye) (****p = 10^-6^), NANOG+dye vs SOX2+dye (****p = 10^-6^), NANOG (no dye) vs SOX2 (no dye) (*p = 0.03)] White diamonds indicate individual field-of-view (FOV) means (n = 179/316/42/82 cells from 12/12/6/6 FOVs for NANOG+dye/SOX2+dye/NANOG (no dye)/SOX2 (no dye)). Black horizontal lines represent group means (mean of FOV means).

**Supplementary Figure 1.**
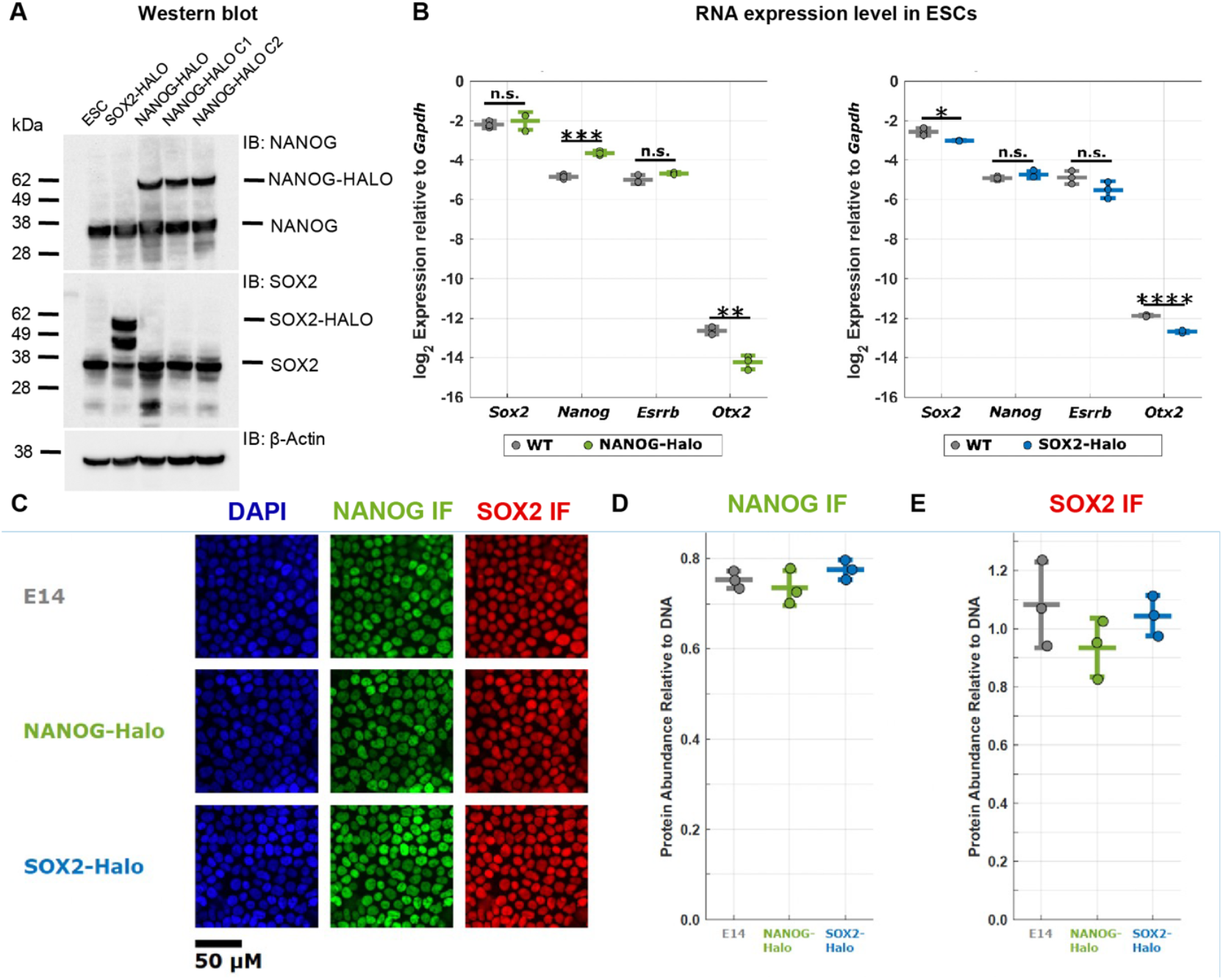
HaloTag-labelled NANOG and SOX2 have similar expression levels and functionality compared to wild-type ESCs. **(A)** Western blot showing that HaloTag cell lines generated are all heterozygous. **(B)** Dot plots comparing RNA levels of naïve pluripotency genes in (left) NANOG-Halo and (right) SOX2-Halo ESCs to wild-type E14 ESCs, as assessed by RT-qPCR of *Sox2, Nanog, Esrrb*, and *Otx2* relative to housekeeping gene *Gapdh* (N=3). Each biological replicate was performed in technical triplicate [Two-tailed unpaired t-test: (left) *Sox2* (p=0.5), *Nanog* (***p=0.00025), *Esrrb* (p=0.07), *Otx2* (**p=0.002); (right) *Sox2* (*p=0.015), *Nanog* (p=0.2), *Esrrb* (p=0.11), *Otx2* (****p=0.00003)]. **(C)** Representative immunofluorescence images showing similar expression of NANOG (green) and SOX2 (red) in wild-type E14, NANOG-Halo and SOX2-Halo ESCs (DAPI, blue) (N=3 for each cell line, with 6 technical replicates for each biological replicate). **(D)** Dot plot of NANOG protein abundance, normalised by DAPI intensity. Each dot represents a biological replicate. [2-way ANOVA: E14 vs NANOG-Halo (p=0.7), E14 vs SOX2-Halo (p=0.6), NANOG-Halo vs SOX2-Halo (p=0.3)]. **(E)** Dot plot of SOX2 protein abundance by cell line, normalised by DAPI intensity. Each dot represents a biological replicate. [2-way ANOVA: E14 vs NANOG-Halo (p=0.3), E14 vs SOX2-Halo (p=0.9), NANOG-Halo vs SOX2-Halo (p=0.5).]

### NANOG has similar chromatin binding ratios to SOX2

Since NANOG is expressed at much lower levels than SOX2, we next wondered whether they had differences in their chromatin binding kinetics. To monitor this, we implemented single-molecule light-field microscopy (SMLFM), a new high-density 3D-SMLM approach we recently established. SMLFM uses a multi-lens array to acquire 7 simultaneous views of the same nucleus taken from different angular perspectives and then extracts the 3D positional trajectories of single molecules through parallax (**Fig.2A, Supp Fig.2**) (Sims et al. 2020; Daly et al. 2024). Although not previously applied to nuclear proteins, SMLFM proved powerful for tracking TFs: its increased z-axis depth of 8 μm captured the entire ESC nucleus (which has a z-axis depth of ∼6 μm) while its ability to acquire localisations at over 10-fold higher density than other PSF-shaping methods allowed us to rapidly capture the spatial distribution of TFs. Since intra-nuclear domains (heterochromatin or mediator condensates) move within the nucleus (Du et al. 2024; Novo et al. 2022), we reasoned this would allow us to better capture spatial distributions. Using SMLFM at 30 ms time resolution, we successfully detected single NANOG/SOX2 molecules at a lateral resolution of 56±6/59±16 nm and an axial resolution of 75±7/79±16 nm (**Supp Fig.3A-B**).

**Figure 2.**
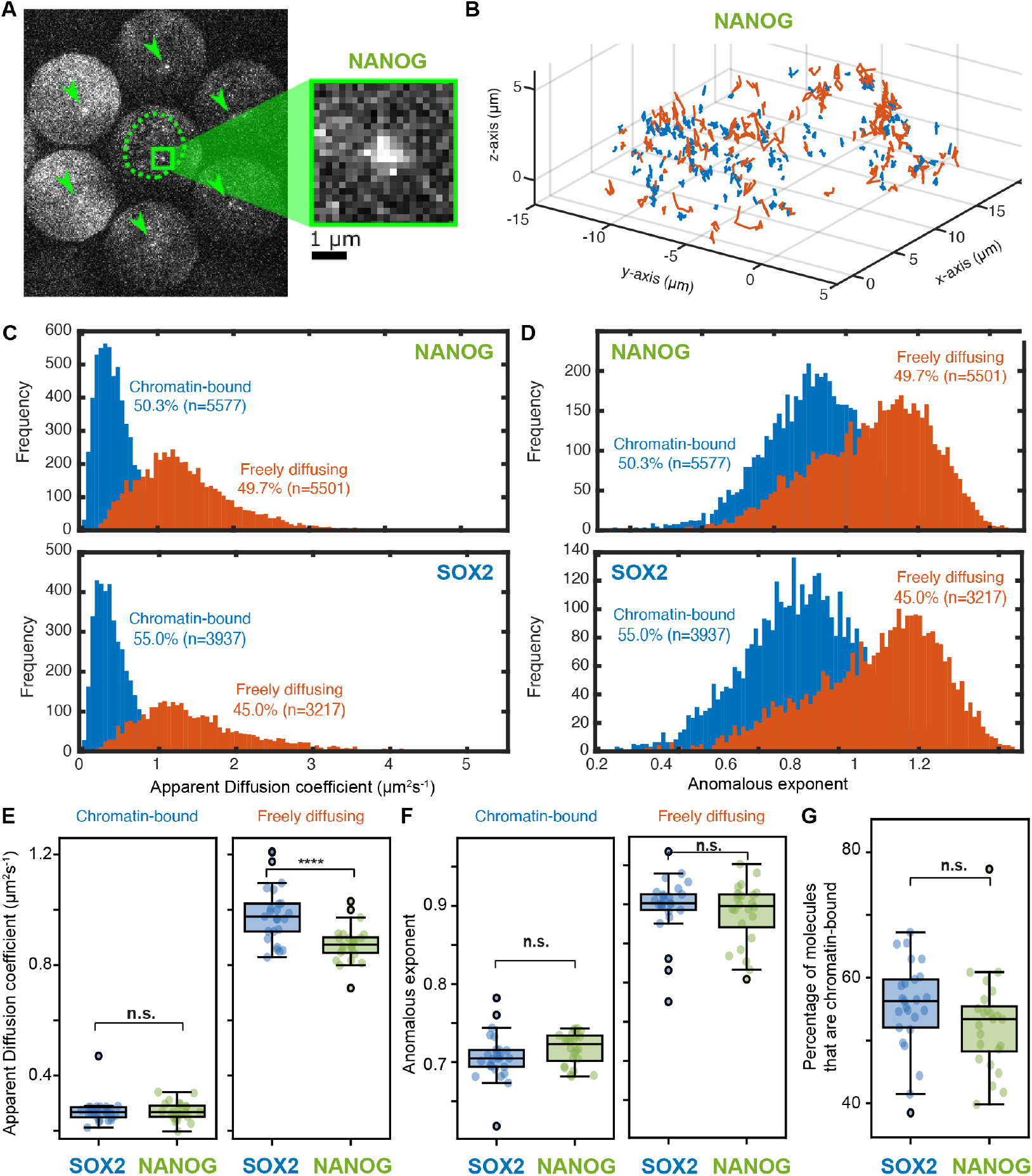
Single-molecule light field microscopy (SMLFM) reveals that NANOG diffuses slower but has a similar chromatin binding ratio to SOX2. **(A)** (Left) Representative SMLFM image showing single PA-JF_646_-HaloTag-tagged NANOG molecules in a live naïve ESC. The nucleus is indicated by a green dotted line based on the sum projection. (Right) Inset showing zoomed in image of a single NANOG molecule. **(B)** Trajectories classified as chromatin-bound (blue) or freely diffusing (orange) for NANOG. **(C)** Histogram showing apparent diffusion coefficient D_app_ of freely diffusing (orange) and chromatin-bound (blue) NANOG and SOX2. **(D)** Histogram showing anomalous exponent of freely diffusing (orange) and chromatin-bound (blue) NANOG and SOX2. **(E)** Boxplot showing D_app_ of chromatin-bound and freely diffusing NANOG/SOX2. [Mann Whitney, p-values: 0.6 (chromatin bound), ****10^-5^ (freely diffusing)] **(F)** Boxplot showing anomalous exponents of chromatin-bound and freely diffusing NANOG/SOX2. [Mann Whitney, p-values: 0.11 (chromatin bound), 0.5 (freely diffusing)] **(G)** Boxplot showing percentage of NANOG and SOX2 molecules classified as chromatin-bound. [unpaired two-tailed t-test, p-value: 0.2] **(B-G)** Dots represent values per FOV (n = 24/25 cells for NANOG/SOX2 imaged from 5 dishes over 2 days).

Having detected single TFs, we next tracked NANOG/SOX2 molecules and used these TF trajectories to analyse the biophysical properties of their freely diffusing and chromatin-bound states. To achieve this, we first calculated high-confidence trajectories of both NANOG and SOX2 (i.e. trajectories that lasted more than 5 frames) (**Supp Fig.3C**) and checked that these trajectories exhibit single-step photobleaching as would be expected from detection of a single molecule (**Supp Fig.3D**). We then applied our published four parameter (4P) algorithm, which uses a sliding window of 11 frames (330 ms) along these trajectories to extract 4 biophysical parameters from single-molecule trajectories (>6800 trajectories/sample were >5 frames) and a Gaussian mixture model to classify and segment trajectories into periods exhibiting either freely diffusing or chromatin-bound behaviour (**Fig.2B-D, Supp Fig.3E-G**) (Basu et al. 2023). Using this method, we discovered that chromatin-bound NANOG and SOX2 had the same apparent diffusion coefficient (D_app_) of 0.27 ± 0.03 μm^2^s^-1^ and 0.27 ± 0.05 μm^2^s^-1^ respectively (**Fig.2C, 2E**). Since the theoretical D_app_ of an immobile molecule is similar at the spatial resolution limit of our microscope, chromatin-bound molecules can simply be regarded as immobile. Meanwhile, freely diffusing SOX2 had a 1.13-fold higher D_app_ than NANOG (0.99 ± 0.11 μm^2^s^-1^ versus 0.88 ± 0.06 μm^2^s^-1^) (p=10^-5^) (**Fig.2C, 2E**), which corresponded to NANOG behaving as if it had a ∼1.5-fold larger hydrodynamic mass than SOX2, according to the Einstein–Stokes relation. Since SOX2 and NANOG are both ∼35 kDa, we concluded that a proportion of NANOG molecules were forming larger complexes, possibly through protein-protein interactions or oligomerisation, consistent with previous observations (Choi et al. 2018; Mullin et al. 2008; Choi et al. 2022).

As a control, we also calculated the anomalous diffusion exponent α from the mean squared displacement (MSD) equation, MSD ∝ time^*α*^, because α is expected to be below 1 for chromatin-bound proteins and close to 1 for freely diffusing proteins. Indeed, freely diffusing NANOG and SOX2 had *α* of 0.90 ± 0.04 and 0.89 ± 0.04, which was higher than that of chromatin-bound NANOG and SOX2, which had *α* of 0.72 ± 0.02 and 0.71 ± 0.03 respectively (**Fig.2D, 2F**). However, freely diffusing NANOG and SOX2 still had *α* values below 1 and so we wondered whether this was because freely diffusing NANOG and SOX2 were not moving by pure Brownian diffusion or whether this was due to a limitation of our 4P algorithm for estimating *α* values. To test this, we simulated Brownian trajectories of NANOG and SOX2 using their average D_app_ values (calculated above) and then used our 4P algorithm to estimate the expected *α* values. This revealed *α* values of 0.96 ± 0.16 and 0.95 ± 0.16, significantly higher than that observed for actual NANOG and SOX2 sub-trajectories (p=10^-74^ and p=10^-209^ respectively) (**Supp Fig.3H**). We therefore concluded that we were accurately classifying chromatin-bound proteins but that our classified freely diffusing trajectories of NANOG and SOX2 were exhibiting more restricted motion than would be expected by pure Brownian diffusion.

To validate these results further, we conducted control SMLFM experiments of immobile HaloTag dye molecules on the coverslip and of cells expressing NLS-tagged HaloTag molecules (**Supp.Fig.4A-B**). As expected, dye molecules on the coverslip did not move in the z-direction and the localisation precision of course exhibited an “apparent” average *D*_app_ and *α* value significantly lower than that of NANOG (p=0.01 and 10^-11^) (**Supp Fig.4C-D**) (this value represents the lower bound limit of detection for the slowest diffusional species). Meanwhile, HaloTag-NLS molecules had average *D*_app_ and *α* values similar to that of NANOG (**Supp Fig.4C-D**). Although Halo-NLS has a lower hydrodynamic mass that NANOG-HALO, this was consistent with previous measurements where adding the positively charged NLS resulted in lower D_app_ values than would be expected from the size of the HALO-NLS (Xiang et al. 2020). These results gave us confidence in our datasets, enabling subsequent analysis.

Using these 30 ms exposure datasets, we then examined how NANOG and SOX2 bind to chromatin. Interestingly, we found that, despite the lower concentration of NANOG, there was no difference in the percentage of chromatin-bound NANOG and SOX2 (53 ± 8% vs 55 ± 7%, p = 0.2) (**Fig.2G**), suggesting that they engage with chromatin to a similar extent.

**Supplementary Figure 2.**
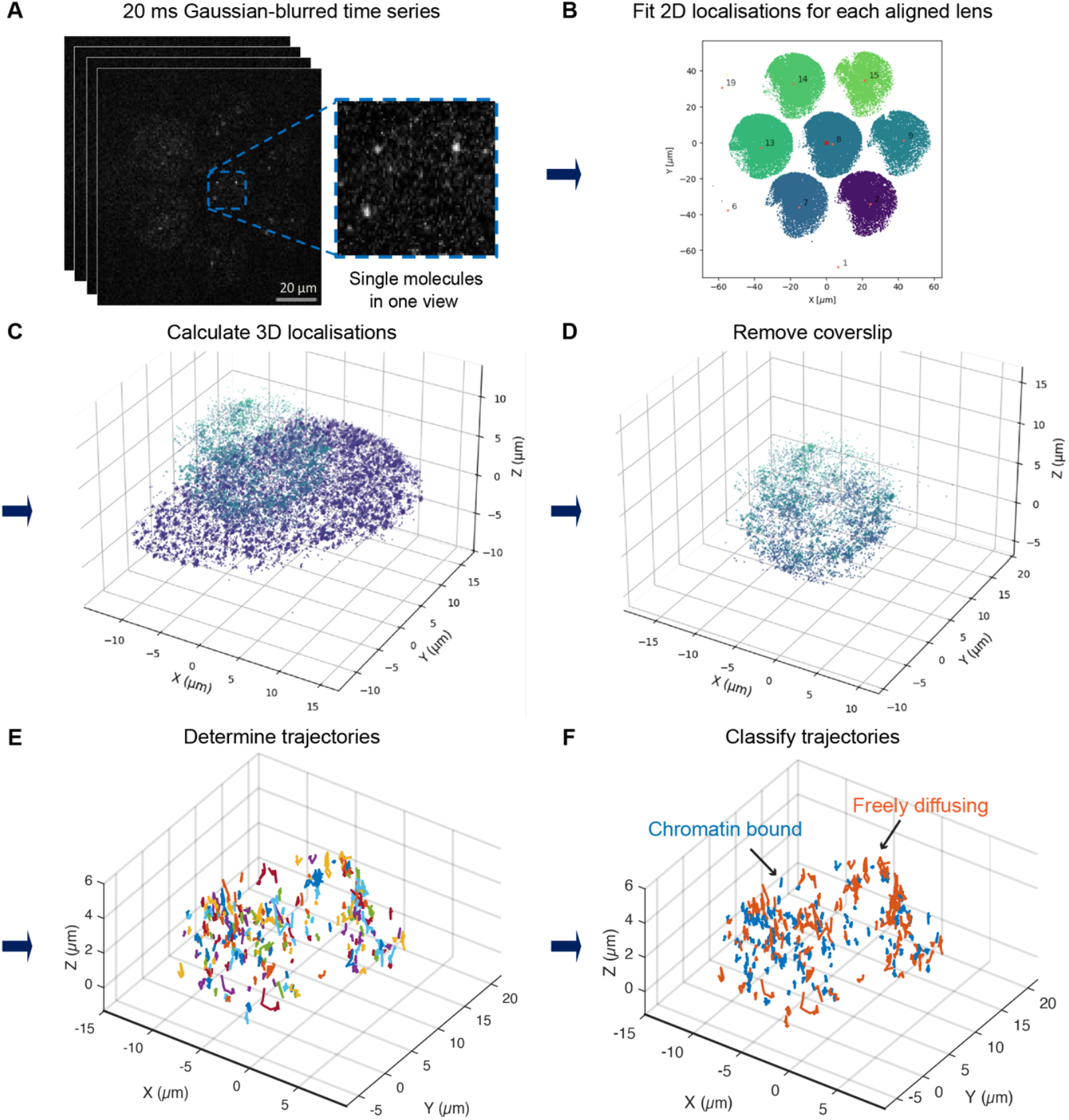
SMLFM image analysis pipeline. **(A)** Representative image from a frame of the video (inset is a zoom of 3 molecules imaged in one of the 7 lenses). **(B)** Images undergo Gaussian fitting to identify 2D localisations that are then assigned through a calibration to the 7 lenses (localisations are coloured by the lens). **(C)** 2D localisations imaged by the different lenses are used to calculate 3D localisations (coloured by z-position). **(D)** The localisations are then curated to remove localisations on the coverslip or those that arise from nearby dirt or cells. **(E)** Localisations are then connected into trajectories (each trajectory in a different colour). **(F)** Trajectories are then classified into sub-trajectories that are either freely diffusing (orange) or chromatin-bound (blue).

**Supplementary Figure 3.**
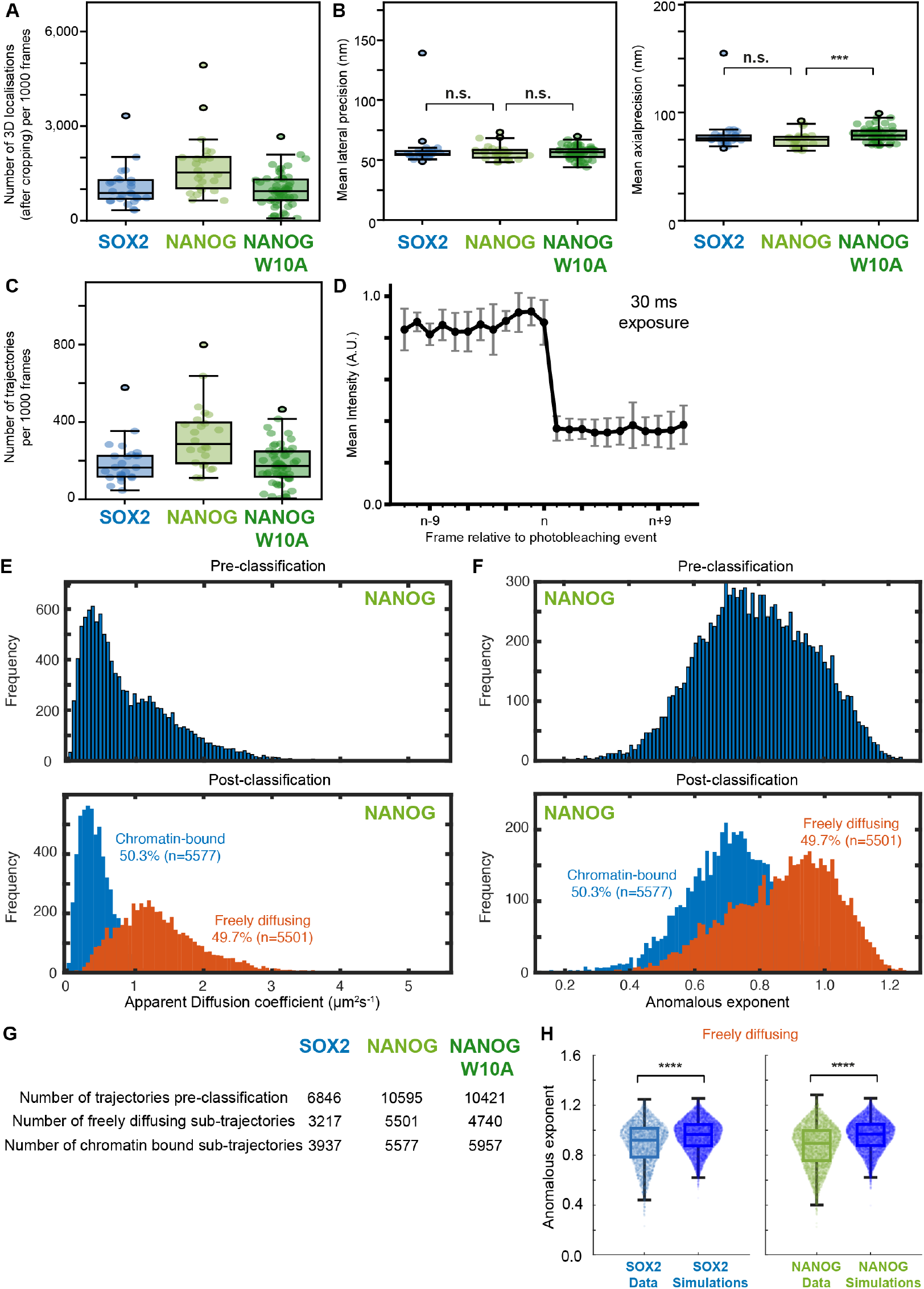
Quality control of 30 ms SMLFM datasets. **(A)** Boxplot showing number of 3D localisations per 1000 frames per cell in each dataset. **(B)** Boxplot showing (left) lateral and (right) axial precision of 30 ms datasets showing no difference in lateral precision [Mann Whitney, p-values: 1.0 (NANOG vs SOX2), 0.6 (NANOG vs NANOG W10A)] but worse axial precision in NANOG W10A. [Mann Whitney, p-values: 0.8 (NANOG vs SOX2), ***10^-3^ (NANOG vs NANOG W10A)] **(C)** Boxplot showing number of tracks per 1000 frames per cell that last for 2 or more frames **(A-C)** Dots represent values per FOV (n = 25/24/56 cells for SOX2/NANOG/NANOG W10A). **(D)** Average single-step photobleaching profile for 30 ms exposure datasets with mean intensity +/− standard deviation shown for 11 frames pre- and post-photobleaching event (n = 10 SOX2 trajectories). **(E)** Histogram showing apparent diffusion coefficient D_app_ of NANOG trajectories (above) pre-classification and (below) post-classification into freely diffusing (orange) or chromatin-bound (blue). **(F)** Histogram showing anomalous exponent of NANOG trajectories (above) pre-classification and (below) post-classification into freely diffusing (orange) or chromatin-bound (blue). **(G)** Table showing the number of total trajectories that are >11 frames, the number of classified freely diffusing trajectories and classified chromatin-bound trajectories for each sample. **(H)** Overlapped scatter and boxplot comparing the anomalous exponent of classified sub-trajectories of freely diffusing SOX2 (blue) and NANOG (green) trajectories to Brownian simulations of the SOX2 and NANOG using their average diffusion coefficients taken from the classified freely diffusing sub-trajectories. [Mann Whitney, p-values: ****10^-74^ (SOX2 freely diffusing trajectories from data vs Brownian simulations), ****10^-209^ (NANOG freely diffusing trajectories from data vs Brownian simulations)]

**Supplementary Figure 4.**
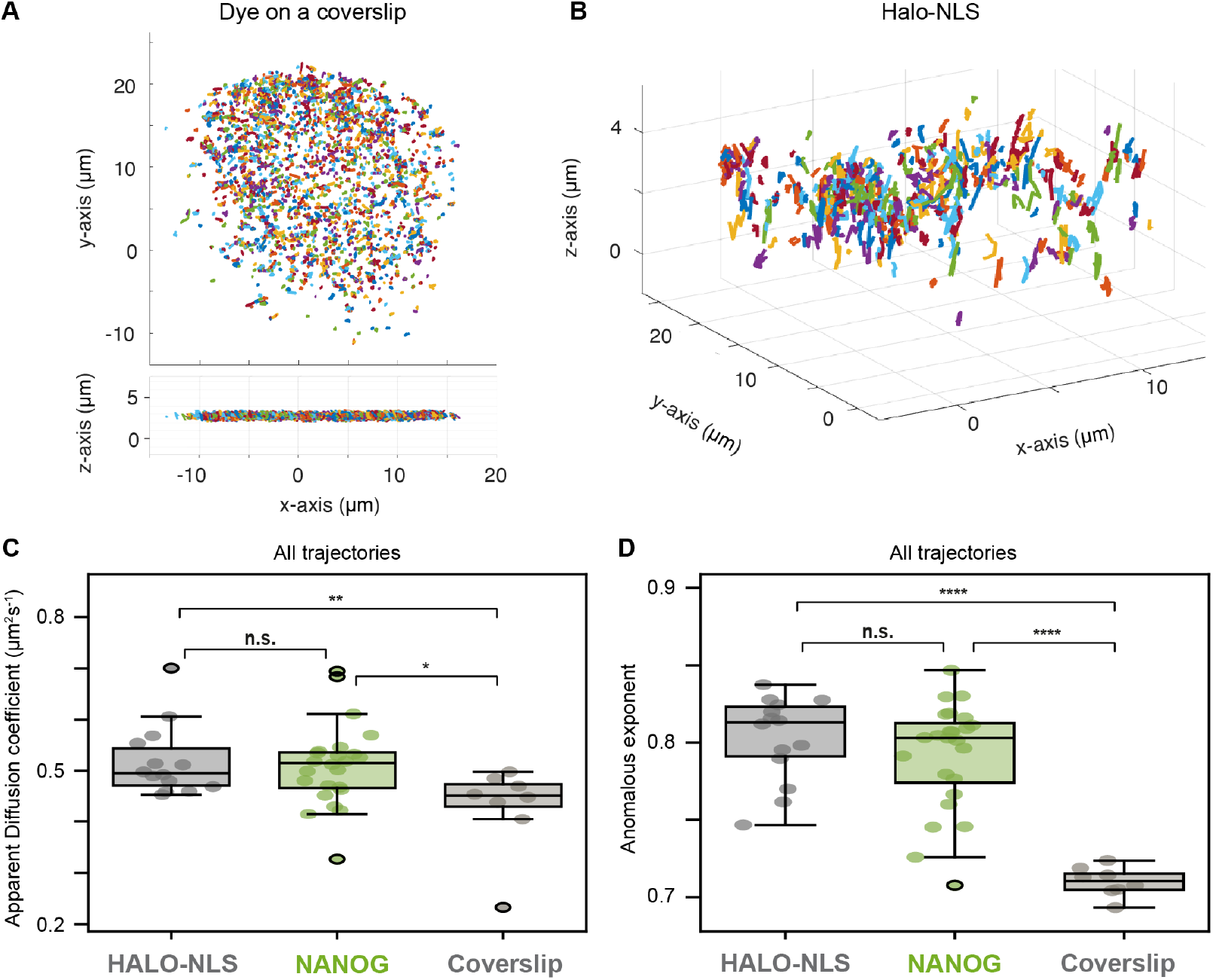
Control datasets of dye on coverslip and HaloTag-NLS. **(A)** Example trajectories of dye on a coverslip shown on an XY and XZ axes to show lack of movement in the z-axis (trajectories labelled in random colours). **(B)** Example trajectories of cells expressing HaloTag-NLS (trajectories labelled in random colours). **(C)** Boxplot showing D_app_ of NANOG trajectories relative to those calculated for HALO-NLS and for dye on a coverslip. [Mann Whitney, p-values: 0.9 (HALO-NLS vs NANOG), **0.008 (Halo-NLS vs dye on coverslip), *0.01 (NANOG vs dye on coverslip)] **(D)** Boxplot showing anomalous exponents of NANOG trajectories relative to those calculated for HALO-NLS and for dye on a coverslip. [t-test, p-values: 0.3 (HALO-NLS vs NANOG), ****10^-9^ (Halo-NLS vs dye on coverslip), ****10^-11^ (NANOG vs dye on coverslip)] **(C-D)** Dots represent values per FOV (n = 14/24/8 FOVs/cells for HALO-NLS/NANOG/coverslip).

### NANOG binds for longer than SOX2

Having observed similar levels of chromatin binding for both NANOG and SOX2, we asked whether there was a difference in how long each TF remains bound to chromatin once they find their DNA binding site i.e. their residence time on chromatin. To measure chromatin residence time, we performed time-lapse SMLFM, based on an approach we have previously described (Gebhardt et al. 2013; Basu et al. 2023). Briefly, it involved detecting only chromatin-bound NANOG/SOX2 molecules by imaging at 200 ms exposures. The longer 200 ms exposure ensured motion blurring of freely diffusing molecules, allowing us to measure how long NANOG/SOX2 are chromatin bound from the number of frames a molecule is detected as chromatin bound before it disappears. However, since disappearance of a molecule may arise not just from chromatin dissociation but also from photobleaching (**Fig.3A**), we collected 4 datasets where we varied the time lapse, τ _tl_, between exposures (0.5s, 2s, 8s, 16s) while keeping the same 200 ms exposure time, τ_int_ (**Fig.3B**). Longer time-lapses reduce photobleaching thereby allowing us to extract true dissociation times along with the photobleaching rate. To further minimise photobleaching, we reduced our laser powers and sacrificed spatial resolution. However, we could still track chromatin-bound NANOG/SOX2 molecules at a lateral resolution of 99±7/91±16 nm and an axial resolution of 138±10/134±19 nm for NANOG/SOX2 molecules (**Supp Fig.5A-C**). As expected for single molecules undergoing chromatin dissociation/photobleaching, we observed molecules disappearing in a single step (**Supp Fig.5D**).

**Figure 3.**
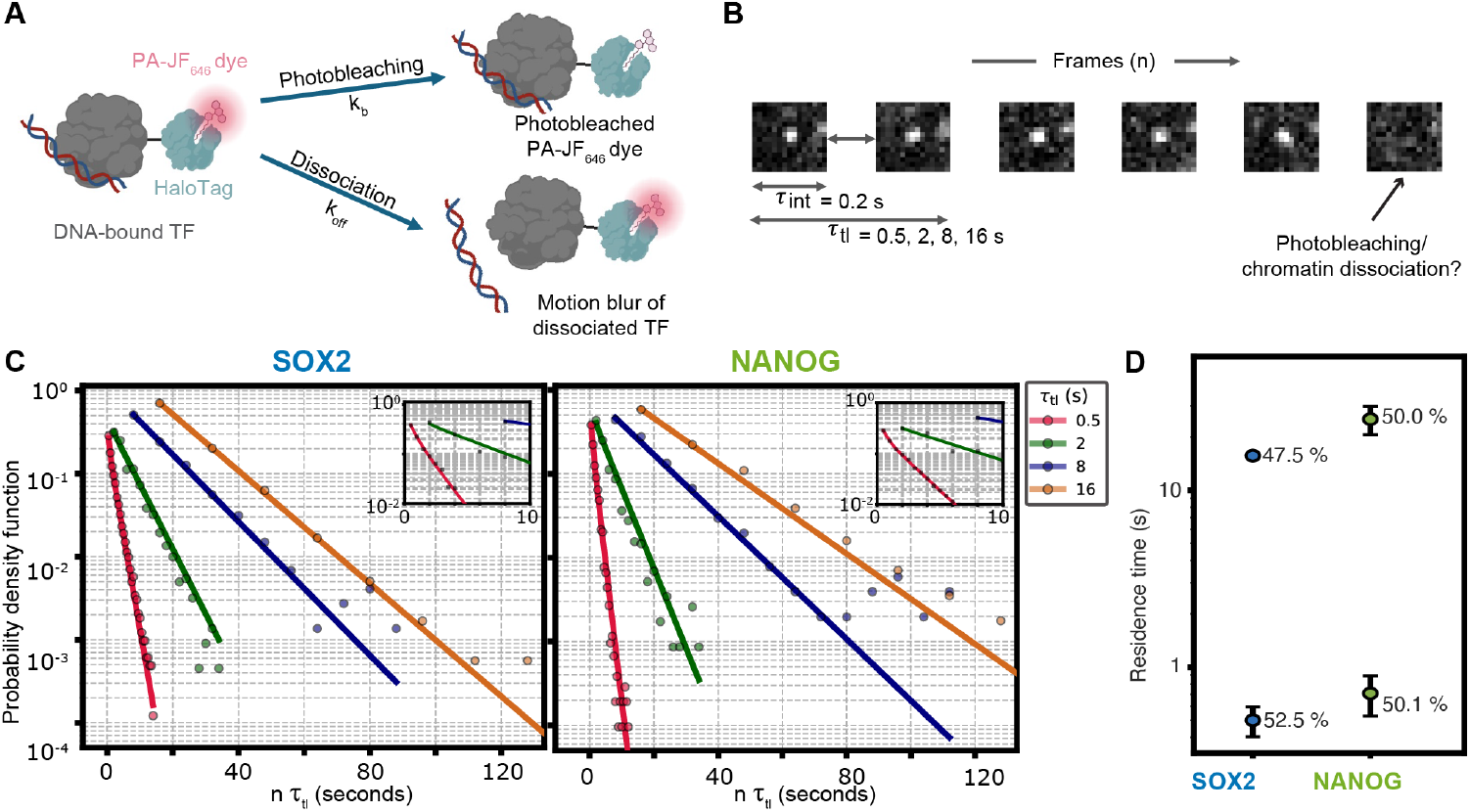
Time-lapse SMLFM reveals that NANOG binds chromatin for longer than SOX2. **(A)** Schematic showing how the measured off rate in time-lapse imaging is determined by two rates: dissociation and photobleaching. **(B)** Schematic showing representative SOX2 molecule from time lapse dataset where the exposure interval, τ_int_, is kept constant at 200 ms exposure while the time lapse between exposures, τ_tl_, is changed (0.5s, 2s, 8s, 16s). Photobleaching rate is correlated to the number of exposures rather than total time, so data collected using variable delays between exposures can be used to disentangle dissociation and photobleaching rates. **(C)** Log plot showing probability density function of (left) SOX2 and (right) NANOG molecules surviving at a given time of imaging where the time is the number of frames multiplied by the time lapse between exposures τ_tl_. Datasets for 0.5s, 2s, 8s and 16s time-lapses (blue, orange, green and red) are fit to a double exponential with a constant photobleaching rate k_b_ and two chromatin dissociation times k_off,1_ and k_off,2_ (for 0.5/2/8/16 s intervals, n = 12/9/4/8 cells for SOX2; n = 10/6/7/9 cells for NANOG). An inset is included zooming into the lower timescales to show the double exponential fitting of the curves. **(D)** Dot plots showing chromatin residence times on a log_10_ axis for NANOG and SOX2, calculated by taking the inverse of k_off_ values extracted from global fitting of graphs in (C).

Having detected and tracked chromatin-bound NANOG/SOX2, we then analysed this data to extract their chromatin residence times. For each time-lapse dataset, we recorded the number of molecules surviving after a given number of frames and fit these values to one, two or three exponential distributions (**Fig.3C, Supp Fig.5E-F**). Bayesian information criterion (BIC) showed that both SOX2 and NANOG datasets fit best to two exponentials, suggesting two modes of chromatin binding (**Supp Fig.5F**). Next, we fit the effective dissociation constant(s) k_eff_ at different timelapses and used them to extract both the photobleaching rate and the true dissociation time using the equation k_eff_ τ_tl_ = k_off_ τ_tl_ + k_b_ τ_int_, where photobleaching was a fixed value related to the exposure time τ_int_. This revealed that SOX2 had two chromatin binding modes with 52.5 ± 2.5 % of molecules having a chromatin residence time of 0.50 ± 0.10 s and 47.5 ± 2.5 % of molecules having a chromatin residence time of 15.7 ± 0.5 s (**Fig.3C, Supp Fig.5E**). This was similar to non-specific and specific binding times previously reported for SOX2 and other TFs (Chen et al. 2014; de Jonge et al. 2022), giving us confidence in our results. NANOG also exhibited two binding modes with 50 ± 6 % of molecules exhibiting each type of binding: one chromatin binding mode had a residence time of 0.71 ± 0.18 s and the other had a residence time of 25 ± 5 s (**Fig.3C, Supp Fig.5E**), slightly higher than that of SOX2. As expected, the photobleaching rate was similar for NANOG and SOX2 – 1.79 ± 0.13 s^-1^ and 1.16 ± 0.05 s^-1^ respectively. We therefore concluded that both NANOG and SOX2 exhibit both non-specific and specific chromatin binding but that NANOG’s specific binding time was longer than that of SOX2, suggesting that it has a larger interaction surface with chromatin, which may be due to its ability to form protein complexes.

**Supplementary Figure 5.**
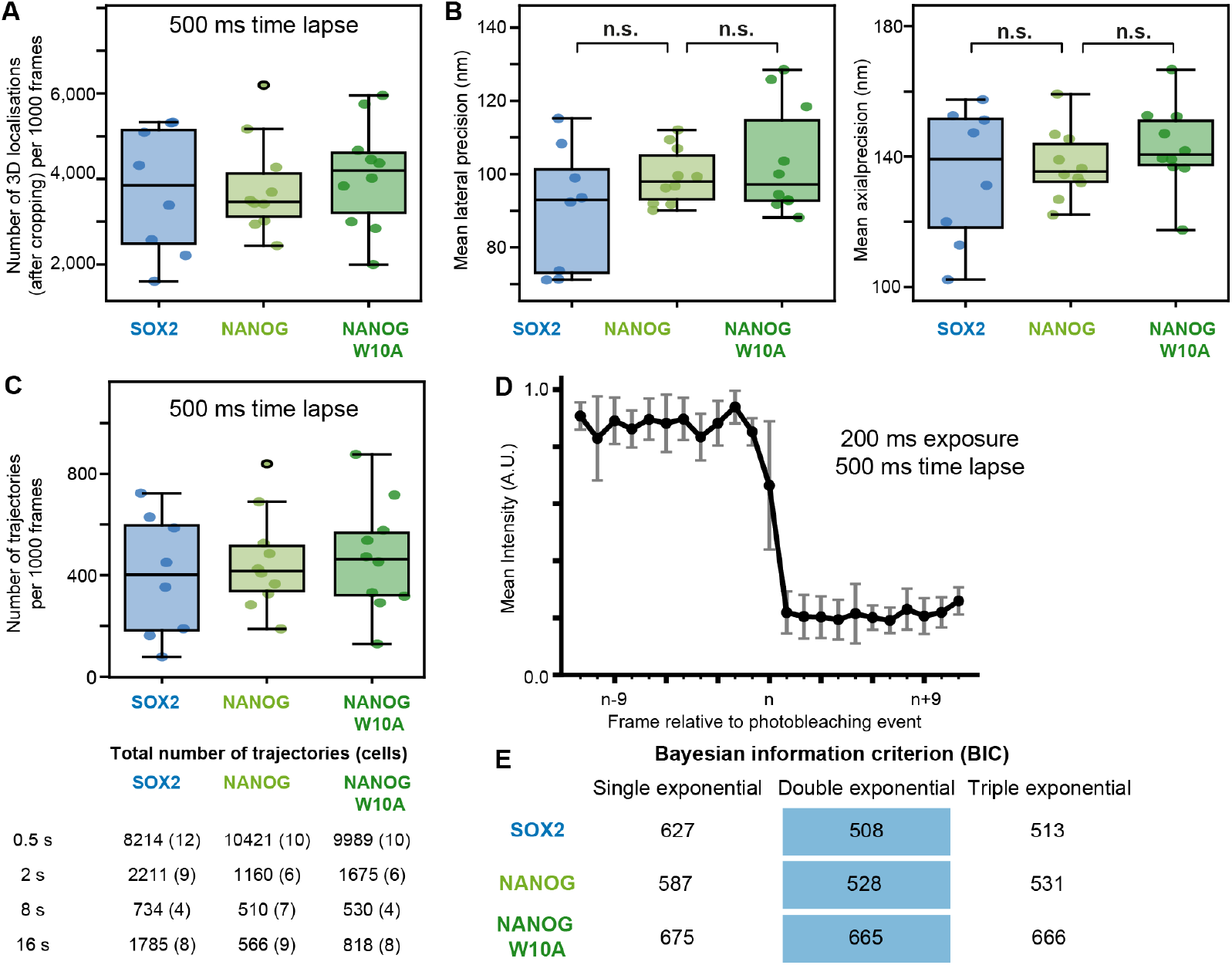
QC of time-lapse SMLFM. **(A)** Boxplot showing number of 3D localisations per 1000 frames per cell in 500 ms time lapse dataset. **(B)** Boxplot showing (left) lateral and (right) axial precision of 200 ms exposure (500 ms time lapse) datasets showing no difference in lateral precision [Mann Whitney, p-values: 0.2 (NANOG vs SOX2), 0.7 (NANOG vs NANOG W10A)] or in axial precision [Mann Whitney, p-values: 0.7 (NANOG vs SOX2), 0.3 (NANOG vs NANOG W10A)]. **(C)** (Top) Boxplot comparing number of SOX2, NANOG or NANOG W10A tracks per 1000 frames per cell that last for 2 or more frames. (Below) Table showing total number of trajectories and cells included in the analysis. **(A-C)** Dots represent values per FOV. **(D)** Average single-step photobleaching profile for 500 ms exposure 500 ms time lapse datasets with mean intensity +/− standard deviation shown for 11 frames pre- and post-photobleaching event (n = 10 trajectories taken randomly from SOX2, NANOG and NANOG W10A datasets). **(E)** Table showing Bayesian information criterion (BIC) for fitting of track lengths to one versus two versus three exponentials to justify chosen fitting in **Fig.3C**.

### NANOG and SOX2 exhibit spatial clustering

Since NANOG and SOX2 have been shown to form phase separated, fibril-like assemblies and aggregates *in vitro* (Krainer et al. 2021; Wang et al. 2021; Choi et al. 2022; Boija et al. 2018), we examined the spatial clustering of NANOG and SOX2 to examine whether or not there was evidence of a phase-separated domain (PSD) within live ESCs. To address this, we established a data-driven spatial analysis approach based on the principle that spatial clustering results in shorter distances between objects within a cluster compared to distances between clusters (**Fig.4A**). 3D Delaunay triangulation was used to compute the distribution of median edge lengths followed by Gaussian mixture modelling (GMM) to determine the number of distinct populations present. This analysis was performed on a per–field of view (FOV) basis. To place the different levels of clustering observed across imaging fields into a broader context, the medians from each FOV Gaussian distribution were pooled, and GMM was again applied to identify clustering levels at the global scale. If more than one population emerged, the population with the smallest median edge length was designated as the clustering population. This global label was then propagated back to all local FOVs. Although we considered using other spatial clustering pipelines (Steindel et al. 2022), we found that most approaches either struggled with cell-to-cell differences in protein density, did not directly identify the clusters themselves (e.g. pair-correlation/nearest-neighbour approaches), required parameter inputs (e.g. DBSCAN) or suffered from edge effects (e.g. Voronoi-based approaches).

**Figure 4.**
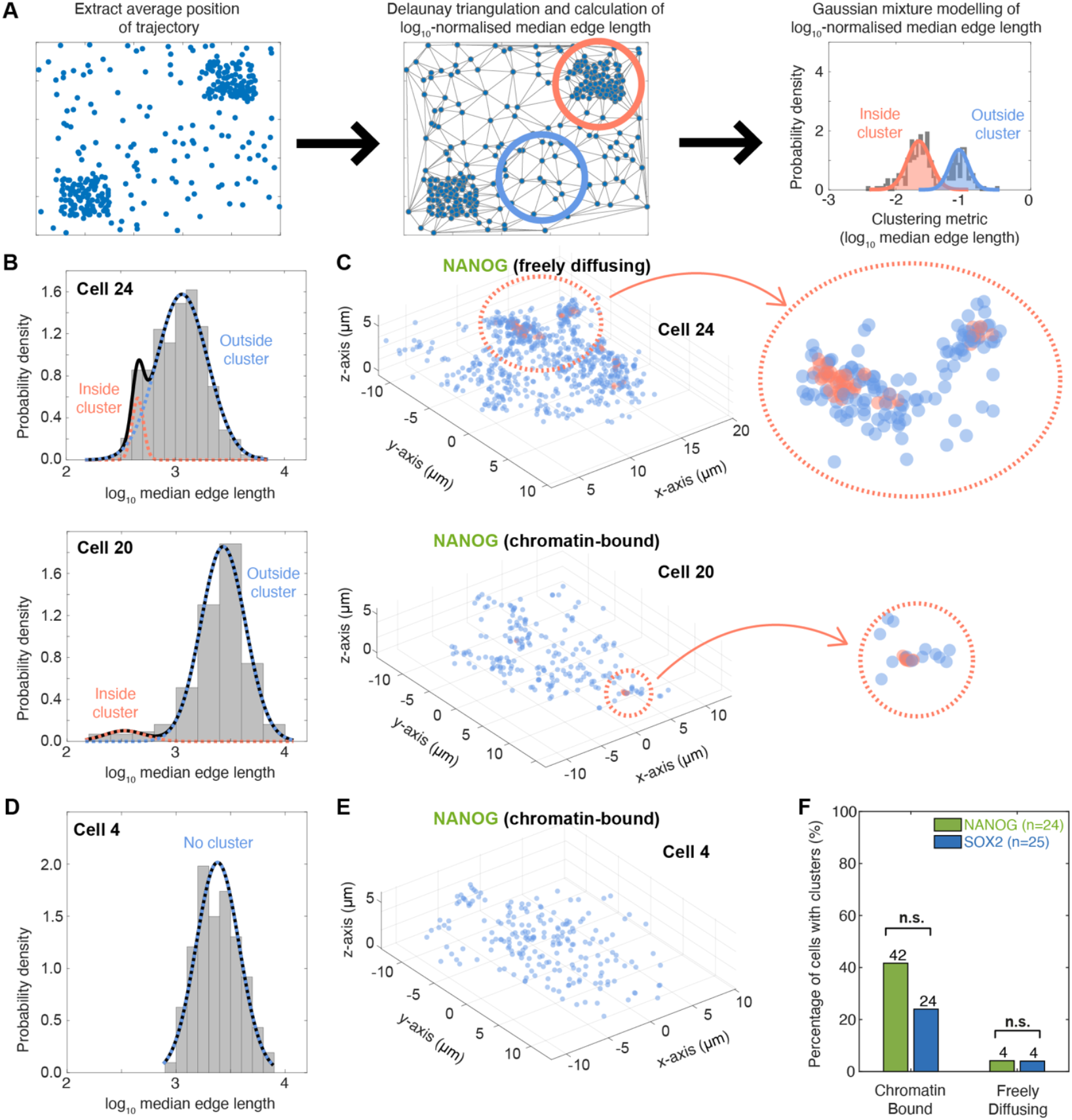
Delaunay-based spatial analysis pipeline reveals clustering of chromatin-bound and freely diffusing NANOG and SOX2. **(A)** Spatial clustering pipeline: (Left) Chromatin-bound and freely diffusing molecules are first identified and their average position determined. (Middle) Delaunay triangulation then connects localisations with edges. The edges are short if there is a cluster (salmon-coloured circle) and longer between clusters (blue circle). (Right) The log_10_-normalised median edge length is then calculated from all particles to produce a distribution that is analysed by Gaussian mixture modelling (GMM) to determine whether there is one population describing these median edge lengths i.e. non-clustered (blue distribution) or whether there is >1 population i.e. clustered (salmon-coloured distribution). **(B)** Histograms of the log_10_-normalised median edge length for the average positions of (top) freely diffusing and (bottom) chromatin-bound NANOG molecules in example cells that show spatial clustering (black line indicates the best fit from the GMM; salmon-coloured line represents the fit to the molecules inside the cluster; blue line represents the fit to the molecules outside the cluster). **(C)** Spatial position of molecules within the cells fit in (B) with salmon-coloured and blue dots representing molecules inside and outside of the cluster respectively. (Right) Inset zooming in on one of the clusters inside the cell. **(D)** Histogram of the log_10_-normalised median edge length for the average positions of chromatin-bound NANOG molecules in an example cell that does not show spatial clustering (black and blue lines indicate the best fit from the GMM). **(E)** Spatial position of molecules within the cell fit in (D) with blue dots representing molecules within the cell. **(F)** Bar plot comparing number of NANOG and SOX2 cells exhibiting clustering using the Delaunay-based spatial clustering pipeline. (n = 25/24 cells for SOX2/NANOG) [Fisher’s exact test, p = 0.2/1 for chromatin-bound/freely diffusing molecules]

To validate our approach, we tested it on simulated single molecules inside of a synthetic nucleus that were either diffusing or that were forming spatial clusters due to an arbitrary interaction force between proteins. As expected freely diffusing proteins exhibited a single distribution of log_10_-normalised median edge distances, where an increase in protein concentration led to a decrease in median edge distances (**Supp Fig.6A-B**). In contrast, the addition of an interaction force between proteins led to spatial clustering and a bimodal distribution of log_10_-normalised median edge distances, as long as protein concentrations were high enough for protein-protein interactions to occur (**Supp Fig.6C-E**). We therefore concluded that this was a powerful way to identify and classify molecules as clustered.

Having established this spatial analysis pipeline, we used it to examine chromatin-bound and freely diffusing TFs separately. This was because, while spatial clustering of freely diffusing molecules may arise from weak protein-protein interactions, clustering of chromatin-bound TFs can arise from repeats of TF binding sites at specific genomic sites or chromatin looping (which brings TF binding sites closer together in 3D). This revealed that both NANOG and SOX2 formed PSDs in some cells but that there was considerable cellular heterogeneity with less than half of cells showing detectable levels of spatial clustering (**Fig. 4B-F, Supp Fig.7A-D**). Moreover, NANOG was as likely to form spatial clusters as SOX2 (p = 0.2/1 for chromatin-bound/freely diffusing proteins) (**Fig. 4F**). Intriguingly, we were considerably more likely to observe clustering of chromatin-bound NANOG/SOX2 than that of freely diffusing proteins (**Fig.4F**), consistent with previous studies showing clustering of chromatin-bound SOX2 proteins (Liu et al. 2014). For the few cells where we observed clustering of freely diffusing molecules, we noted that, while freely diffusing molecules within a cluster exhibited no change in D_app_ (p=0.2), they had lower anomalous exponent (p = 0.03), indicative of more restrictive motion (**Supp Fig.7E**). We concluded that both NANOG and SOX2 can form PSDs in principle and that these PSDs were characterised by spatial clustering of chromatin-bound and sometimes freely diffusing NANOG and SOX2.

**Supplementary Figure 6.**
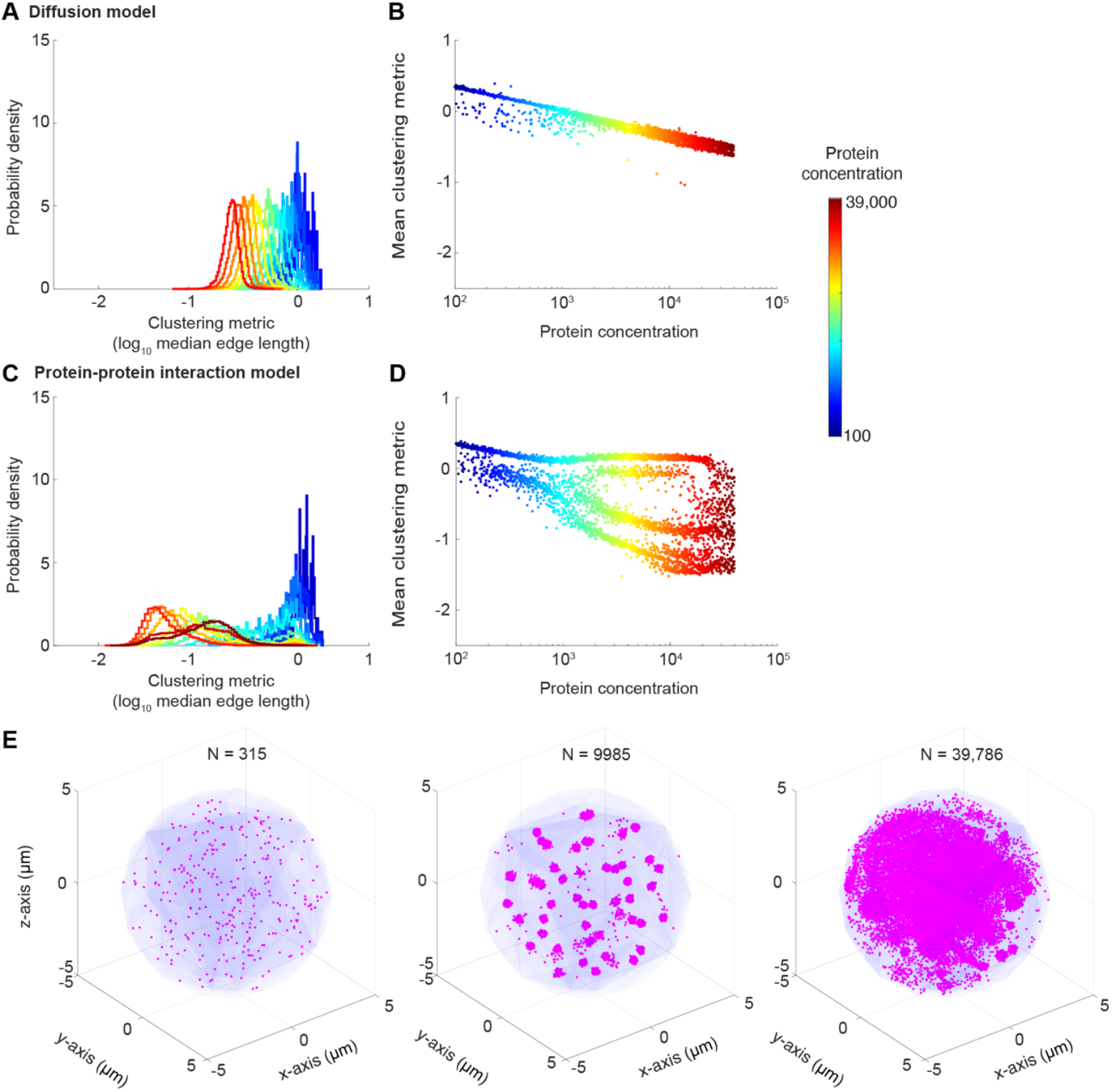
Validation of Delaunay-based clustering pipeline. **(A)** log_10_-normalised median edge lengths extracted from the end position of proteins simulated using a diffusion model showed no change in the clustering metric distribution with increased protein concentration leading to a shift in the distribution to lower values. **(B)** Gaussian mixture modelling (GMM) of distributions in (A) showed no distinct sub-populations. **(C)** log_10_-normalised median edge lengths extracted from the end position of proteins simulated using a protein-protein interaction model showed a shift to a bimodal distribution at intermediate concentrations because protein-protein interactions are rare at low concentrations and because all proteins end up in one phase-separated domain within the nucleus at high concentrations. **(D)** GMM of distributions in (C) showed a separation of sub-populations at intermediate protein concentrations. **(E)** Example realisations of the protein-protein interaction model at different protein concentrations. **(A-E)** Simulations were run for 1300 particle concentrations from 10^2^ to 10^4.6^, increasing linearly in the exponent. Total simulation time was 2 s. Time step was 0.02 s. Model parameters were set as D = 0.1 μm^2^s^−1^, spherical potential well radius for protein-protein interactions was 1 μm, and potential well power was 0.075.

**Supplementary Figure 7.**
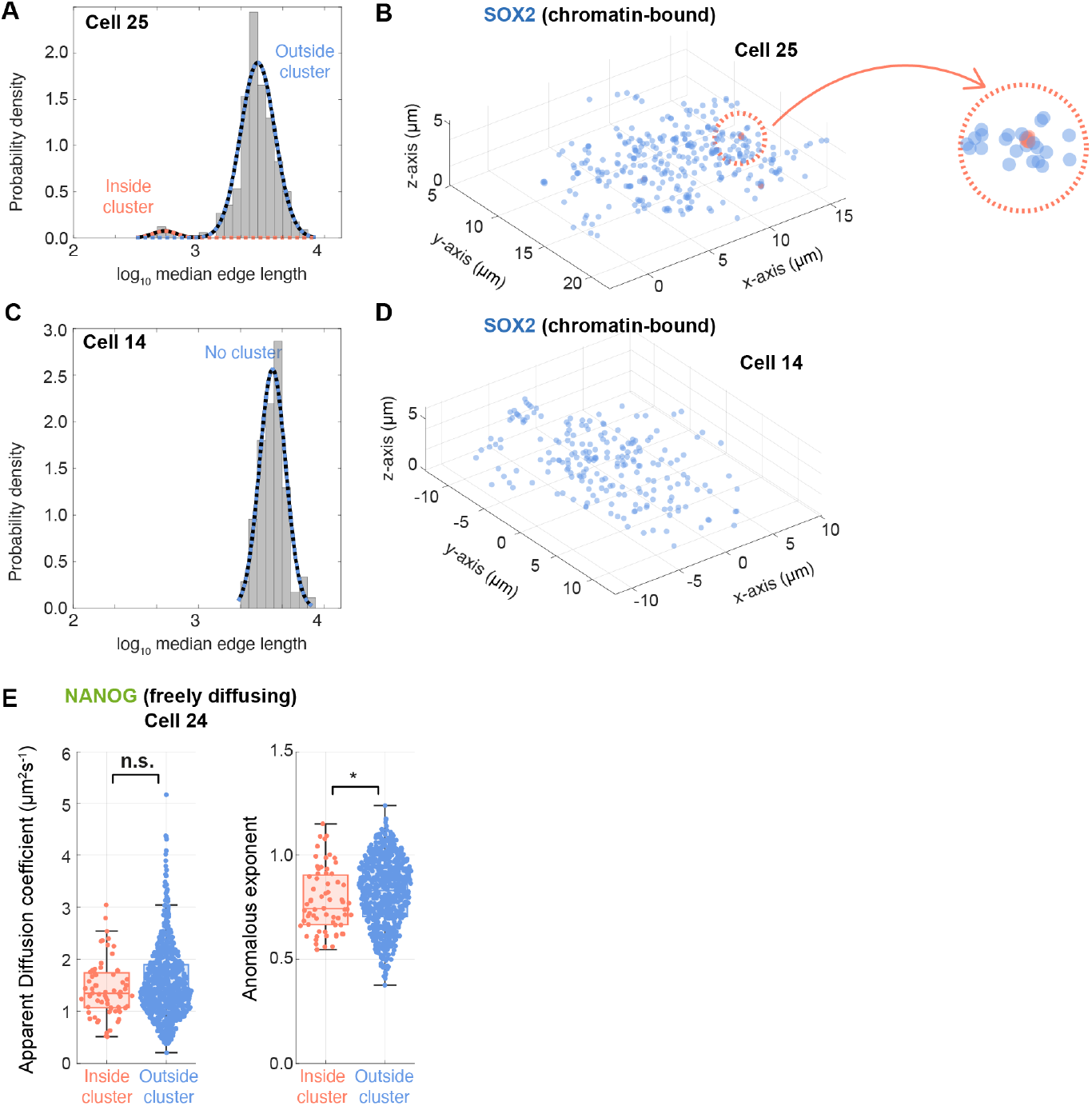
SOX2 exhibits spatial clustering and comparison of freely diffusing NANOG inside and outside of spatial clusters. **(A)** Histograms of the log_10_-normalised median edge length for the average positions of chromatin-bound SOX2 molecules in example cell that shows spatial clustering (black line indicates the best fit from the GMM; salmon-coloured line represents the fit to the molecules inside the cluster; blue line represents the fit to the molecules outside the cluster). **(B)** Spatial position of molecules within the cell fit in (A) with salmon-coloured and blue dots representing molecules inside and outside of the cluster respectively. (Right) Inset zooming in on one of the clusters inside the cell. **(C)** Histogram of the log_10_-normalised median edge length for the average positions of chromatin-bound SOX2 molecules in an example cell that does not show spatial clustering (black and blue lines indicate the best fit from the GMM). **(D)** Spatial position of molecules within the cell fit in (C) with blue dots representing molecules within the cell. **(E)** Boxplot comparing (left) D_app_ and (right) anomalous exponents of freely diffusing NANOG inside and outside of clusters identified in cell 24 [Mann Whitney, p-values: 0.2 (D_app_), *0.03 (anomalous exponents)]

### SOX2 forms larger intra-nuclear phase-separated domains (PSDs) than NANOG

Having shown that chromatin-bound NANOG and SOX2 molecules were spatially clustered in some cells and that there was also some evidence of spatial clusters containing freely diffusing molecules (**Fig.4B-C**), we wondered whether freely diffusing NANOG/SOX2 molecules were within a phase-separated domain (PSD) that restricts the diffusion of NANOG/SOX2. This would account for the fact that freely diffusing NANOG and SOX2 had *α* values that were lower than would be expected for molecules undergoing Brownian diffusion (**Supp Fig.3H**) and for the fact that freely diffusing molecules had lower anomalous exponents within a cluster (**Supp Fig.7E**). If NANOG/SOX2 were within PSDs, this would mean that NANOG, despite its low concentration, would have a smaller region of the nucleus to explore to find its binding sites, considerably decreasing chromatin association times within these PSDs. However, it was a challenge to determine how NANOG/SOX2 diffuse within PSDs and to quantify the size of these PSDs. If a sufficiently large number of trajectories had been available, PSDs could have been identified as regions where freely diffusing trajectories within the same cell were confined. However, since NANOG molecules within these regions were limited in number, we needed to develop an approach that compares the spatial exploration of trajectories across different cells based on the principle that molecules move differently over time when freely diffusing versus when they are confined to a PSD (hereafter referred to as ‘domain-confined’).

To address this challenge, we established an approach that uses the convex hull associated with a trajectory developing over time, specifically one that analyses how the perimeter of this convex hull changes. The convex hull describes the smallest convex shape that encloses all points within the trajectory over a given time period and, theoretically, the perimeter of this convex hull increases differently if a molecule is diffusing within a confined PSD versus if it is diffusing in an unconfined manner (**Fig.5A**). We reasoned that we could use this difference to distinguish molecules within a PSD from those that were not. We therefore established a classification approach based on a hybrid machine learning algorithm that uses time-series clustering coupled to convex hull-perimeter dynamics (**Supp Fig.8**). The pipeline took input trajectories and computed both the time-dependent convex hull and its perimeter (**Supp Fig.8A-C**). To evaluate the similarity between trajectories that could be confined at different places, we then used a k-means clustering framework (Pelleg and Moore 1999; Lloyd 1982) where the distance between two curves was computed from Dynamic Time Warping (DTW) (Sakoe and Chiba 1978) (**Supp Fig.8D**). The DTW-K-means algorithm generated three outputs: an ensemble of clustered perimeter curves (**Supp Fig.8E**), the percentage of molecules that were domain-confined/unconfined and the labelled original trajectories classified as domain-confined or unconfined (**Supp Fig.8F**). Finally, from the domain-confined trajectories, we extracted the radius 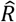 of confinement of the underlying PSD by fitting the average perimeter vs time curve to the theoretical formula (**Supp Fig.8G-I**) (De Bruyne et al. 2022). This radius provides an estimate of the average observed PSD size and, although the PSD size is likely to be variable at the single-cell level, the average PSD served as a parameter that allowed relative comparisons between datasets.

**Figure 5:**
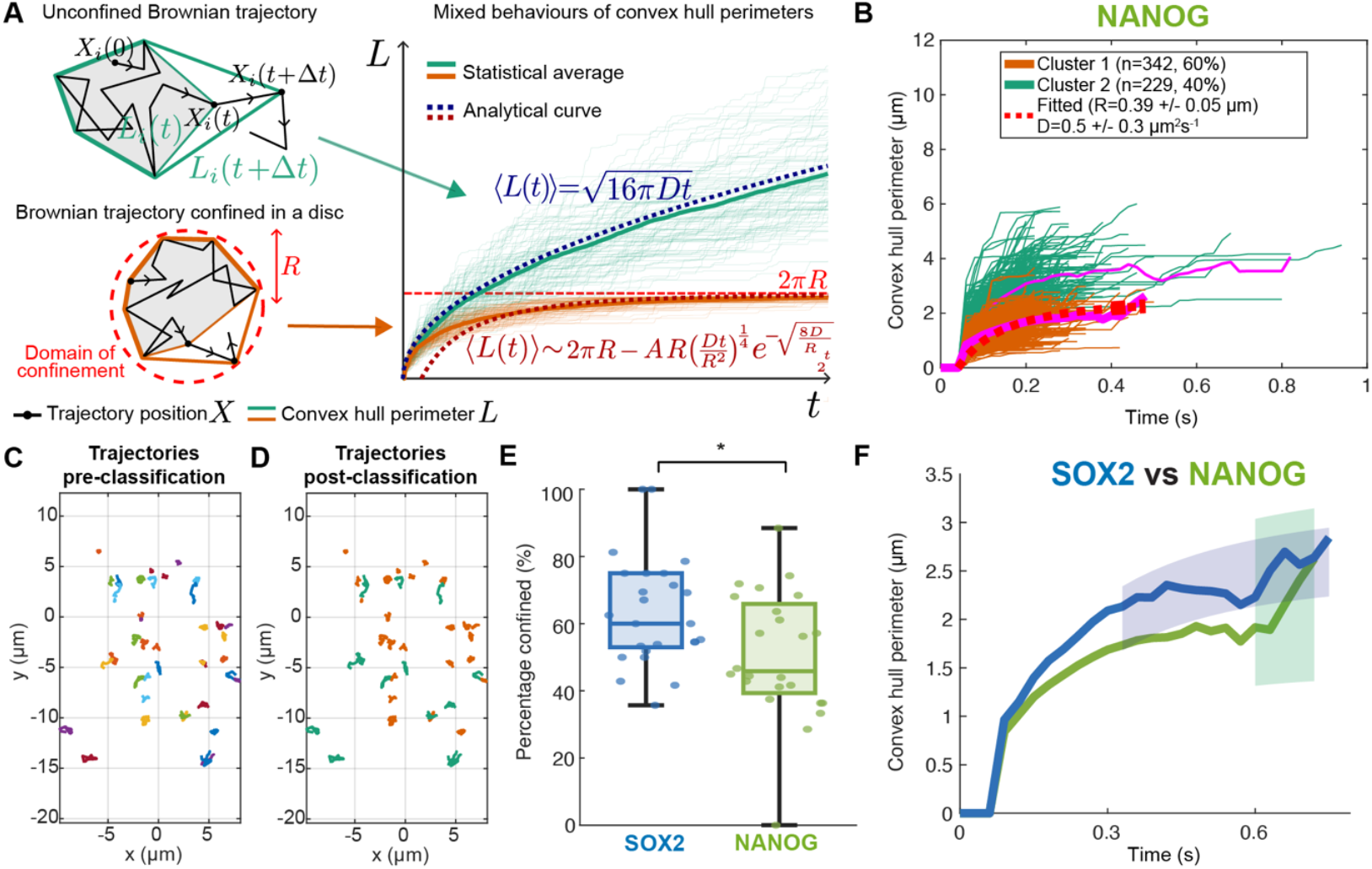
Pipeline for computing convex hull perimeters over time reveals larger phase-separated domains for SOX2 than NANOG. **(A)** Growth of the convex hull differs for a free Brownian trajectory versus one that is confined to a reflective disc. For a Brownian trajectory with positions *X*(*t*), the perimeter *L* of convex hull increases as a function of time *t*. In contrast, the growth of the convex hull is limited for a Brownian trajectory confined within a reflective disc representing a PSD. Convex hull perimeters of open and domain-confined Brownian trajectories show distinct behaviours. The average perimeter of open Brownian trajectories grows as 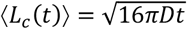, where *D* is the diffusion coefficient (Casalis, Letac, and Massam 1993); whereas, via asymptotic analysis the mean convex hull perimeter of the Brownian trajectories confined in a disc converges slowly to the disc’s circumference (De Bruyne et al. 2022): 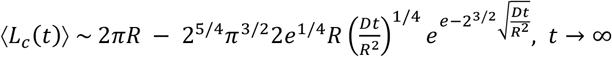. **(B)** The convex hull analysis pipeline clusters freely diffusing NANOG into domain-confined (orange) and unconfined (green) populations. Pink lines depict average values from classified trajectories while red dotted line represents fitted curve for the statistical average curve of domain-confined trajectories. (n = 24 cells) **(C)** Example freely diffusing NANOG trajectories from a cell pre-classification with each trajectory coloured randomly. **(D)** Example freely diffusing NANOG trajectories post-classification: green trajectories are those classified as unconfined and orange trajectories are domain-confined. **(E)** Boxplot showing greater percentage of domain-confined molecules at the single-cell level for SOX2 (blue) than for NANOG (green) (n = 25/24 cells for SOX2/NANOG) [Wilcoxon rank sum test, *p=0.01]. **(F)** Comparing the mean and 95% confidence interval of the fitted radii shows a smaller and greater range of PSD radii for NANOG (green) than for SOX2 (blue). (n = 25/24 cells for SOX2/NANOG)

To validate our new approach, we tested it on synthetic data (Brownian simulation trajectories) and showed that we could recover trajectories confined in a small disc from those that were unconfined at a classification accuracy of 83 ± 6 % and a radius fit accuracy of 93 ± 3%, where most of the error originated from ground-truth unconfined curves being mis-classified as domain-confined (**Supp Fig.9A-F**). We then varied the size of a disc to determine how well the algorithm could estimate the confinement radius of the disc. This revealed that our algorithm was extremely reliable at outputting the confinement radius but that there was an upper limit to the size of PSDs that could be detected (**Supp Fig.9G-I**). This upper limit was dependent on the diffusion speed of the molecules and so, for molecules like freely diffusing NANOG/SOX2 that exhibit a D_app_ of ∼1-1.5 μm^2^s^-1^, we observed a decrease in the accuracy of our algorithm at PSDs greater than 1.4 μm (**Supp Fig.9H-I**).

Having established and validated our approach, we then applied it to freely diffusing NANOG and SOX2 trajectories (**Fig.5B**). This revealed that they can both be classified as domain-confined and unconfined (**Fig.5C-D**) but that SOX2 molecules were more likely to be part of PSDs than NANOG (p = 0.01) with 52 ± 26 % of NANOG and 66 ± 18 % of SOX2 molecules found within PSDs (**Fig.5E**). Intriguingly, we now saw domain-confined molecules in most cells, even those where we had not previously observed clustering, suggesting that NANOG and SOX2 are likely part of PSDs that are predominantly enriched for other molecules and so they are at too low levels within a specific PSD to exhibit spatial clustering. Comparing these classified trajectories to simulations of freely diffusing and domain-confined trajectories further validated our classifications (**Supp Fig.9J-K**). Moreover, while NANOG/SOX2 trajectories could not be described purely by unconfined diffusion, HALO-NLS trajectories fit perfectly to simulated trajectories of unconfined diffusion as expected (**Supp Fig.9L**). Since a proportion of both NANOG and SOX2 were moving within a confined PSD, we next asked whether there were any differences in the size of NANOG-versus SOX2-containing PSDs. Fitting of domain-confined trajectories estimated the radius of confinement of these PSDs to be 0.39 ± 0.05 µm for NANOG, smaller than the 0.447 ± 0.017 µm observed for SOX2 (**Fig.5D**). However, the fitting also revealed a much larger range of possible radii for NANOG-containing PSDs with SOX2-containing PSDs being at the upper end of this range. Since these values were all within the detection limit of our algorithm, we concluded (1) that a relatively large proportion of NANOG and SOX2 were trapped in PSDs of ∼0.4 µm; (2) that SOX2 has a greater likelihood than NANOG of being within PSDs and of forming larger PSDs.

**Supplementary Figure 8:**
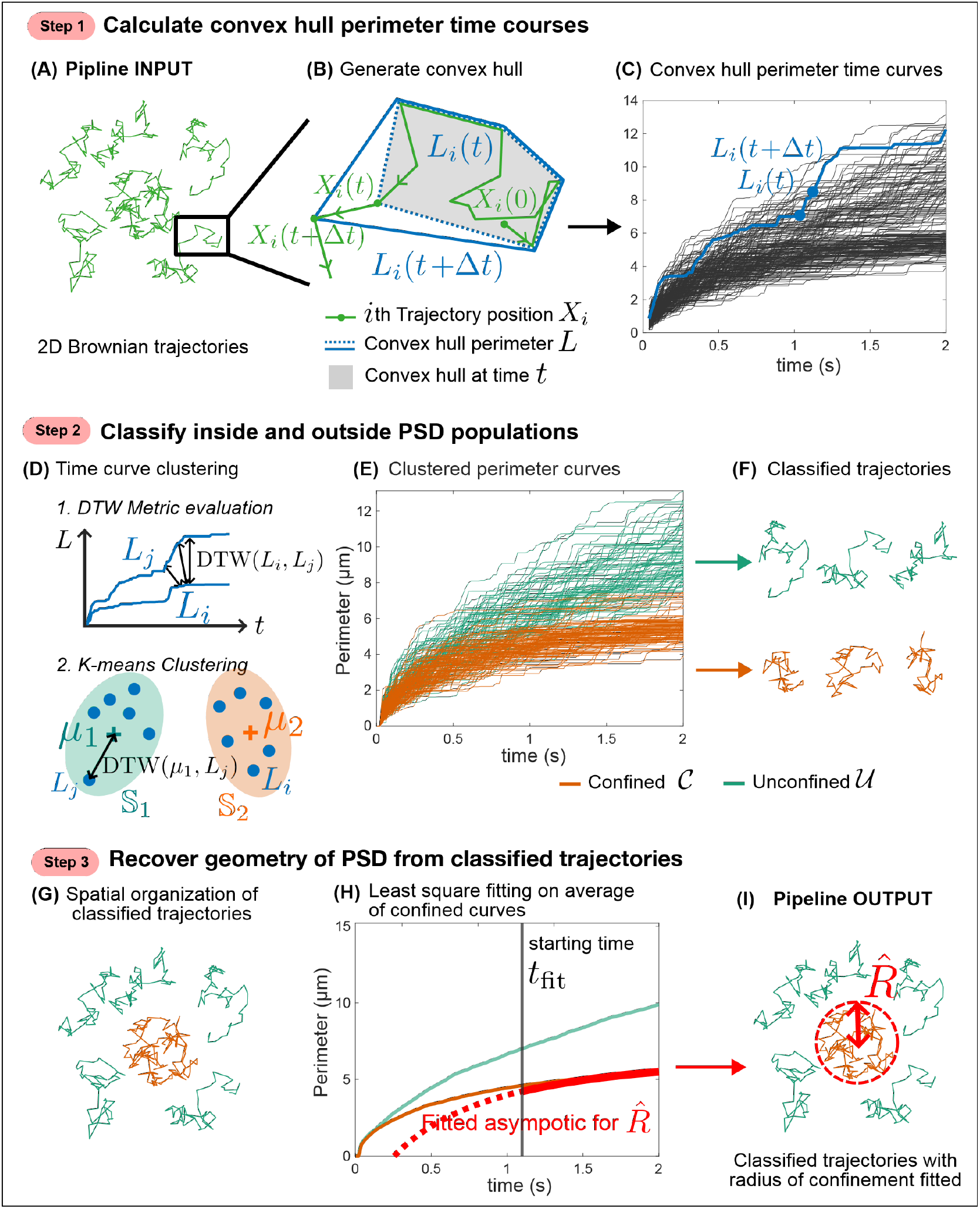
Convex hull analysis pipeline classifies proportion of molecules within phase-separated domains and radius of confinement of the domains. Our approach developed to classify trajectories into domain-confined and unconfined populations; the data shown is from simulating Brownian particles inside and outside a reflective circular boundary. Step 1, **(A)** the pipeline takes 2D Brownian trajectories as input, and **(B)** for each trajectory generates convex hulls at all time steps. The perimeters of the convex hulls grow as a function of time. **(C)** The perimeter curves of all trajectories are collected. Step 2, **(D)** time curves can be clustered based on dynamic time warping (DTW)-K-means method. **(E)** Collected perimeter curves and **(F)** the corresponding trajectories are clustered into either domain-confined (C, orange) or unconfined (U, green). Step 3, **(G)** Classified inside trajectories may spatially co-occur at potential domain of confinement. To extract the radius *R* of the assumed circular PSD, **(H)** the statistical average curve ⟨*L*(*t*)⟩ of the inside population is fitted to the asymptotic equation for 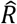. **(I)** The pipeline outputs the fitted radius of confinement 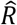 and the classified trajectories.

**Supplementary Figure 9:**
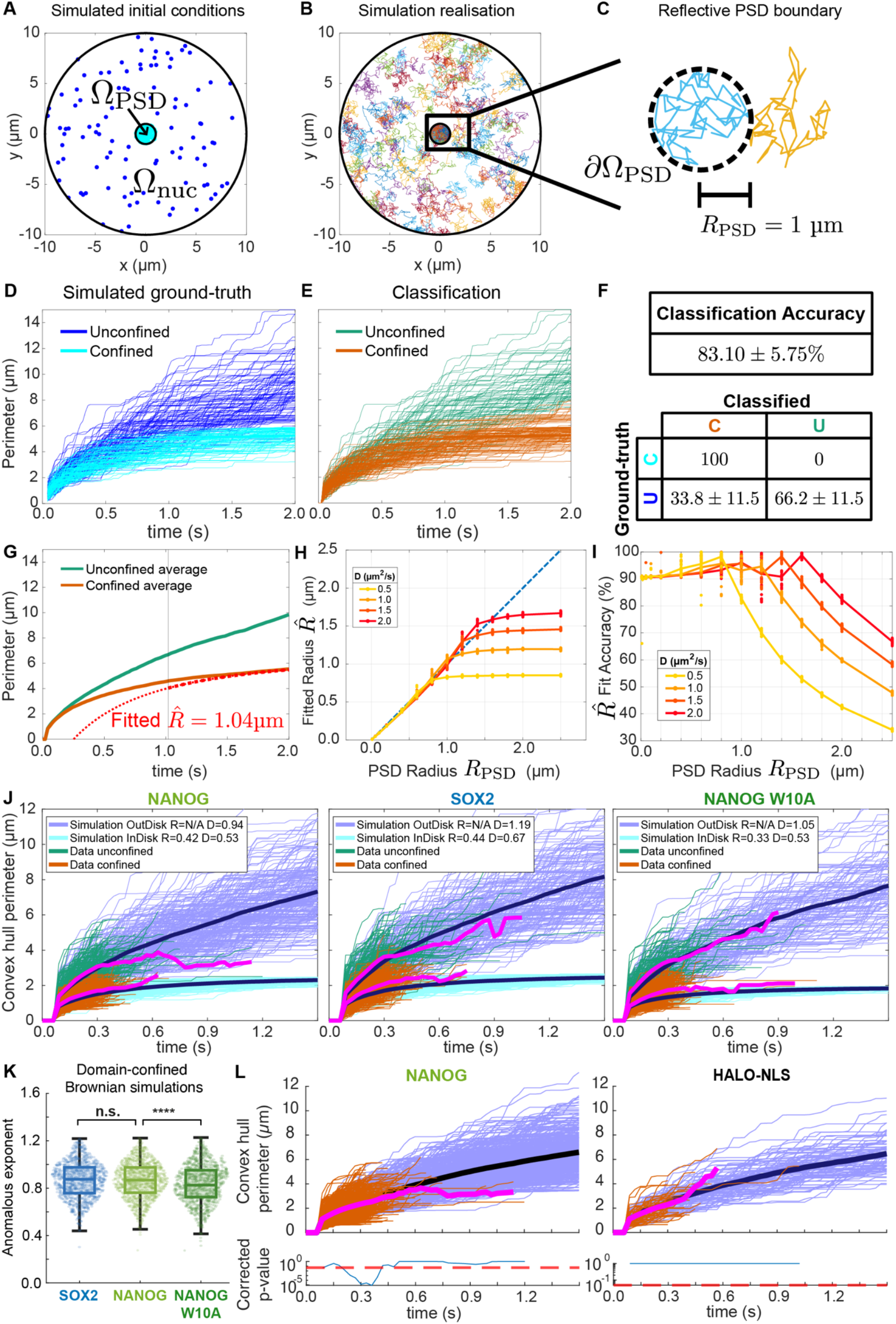
Validation of convex hull pipeline using simulation data. **(A)** As an example, we simulated an inner disc of *R* = 1µm to represent the phase-separated domain (PSD) and an outer disc of *R*_nuc =_ 10µm to represent the nuclear membrane, with 100 particles both inside (light blue) and outside (dark blue) the PSD. **(B)** An example realization shows the trajectories are Brownian of 2-second total length with Δ*t* = 0.02 seconds. We ran 200 realizations. **(C)** Zoom-in on the PSD boundary *∂*Ω_PSD_ shows its reflective property for both inside and outside. **(D)** Ground-truth *L*(*t*) curves computed from the simulation. **(E)** The pipeline clusters unlabeled curves into inside and outside populations. **(F)** The average clustering accuracy (±sd) is 83.10 ± 5.75%. The confusion matrix suggests mis-classification of outside curves to inside. **(G)** Estimation of the PSD radius 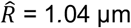 from fitting the ⟨*L*(*t*)⟩ curve of the inside population. **(H)** and **(I)** There exists a threshold of the largest 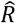 that can be accurately estimated compared to the set PSD radius *R*, which depends on diffusion coefficient *D* of the particles. **(J)** Ground-truth *L*(*t*) curves from simulations of unconfined and domain-confined trajectories compared to classified trajectories from NANOG (left) and SOX2 (middle) and NANOG W10A (right) datasets. Real and simulated trajectories shown in green and dark blue for unconfined and orange and light blue for domain-confined. Average of real and simulated trajectories shown as pink and black lines respectively (n = 25/24/56 cells for SOX2/NANOG/NANOG W10A). **(K)** Overlapped scatter and boxplot comparing the anomalous exponent of simulated domain-confined and unconfined trajectories of SOX2 (blue), NANOG-WT (light green), and NANOG-W10A (dark green). For each TF, 1000 trajectories were simulated using the average diffusion coefficient calculated from the classified freely diffusing trajectories and the domain size estimated by our convex hull pipeline. [Mann Whitney, p-values: 0.9 (SOX2 vs NANOG WT), ****10^-5^ (NANOG WT vs NANOG W10A)] **(L)** Ground-truth *L*(*t*) curves from simulations of unconfined trajectories compared to unclassified trajectories from NANOG (left) and HALO-NLS (right) with p-values calculated below to compare values, showing that HALO-NLS can be described as unconfined but that NANOG trajectories cannot. (n = 24/14 FOVs/cells for NANOG/HALO-NLS).

### NANOG’s oligomerisation domain increases the stability and robustness of chromatin binding at the single-cell level

Having observed that NANOG has a unique ability to bind for longer than SOX2 and to exhibit spatial clustering on DNA, we wondered whether these properties were linked to NANOG’s interactions with SOX2 and/or itself through its WR domain (Choi et al. 2018; Mullin et al. 2008; Choi et al. 2022; Gagliardi et al. 2013). To address this, we generated knock-in ESCs expressing NANOG-Halo with a W10A mutation where the 10 tryptophan (W) residues required for these protein-protein interactions were mutated to alanine (A) (**Supp Fig.10A**). Immunofluorescence experiments confirmed that these cells retained naïve pluripotency with similar levels of NANOG/SOX2 (**Supp Fig.10B-D**). Western blots confirmed that knock-in ESCs were heterozygous for the mutation (**Supp Fig.1A**).

We first tested whether the W10A mutation would impact the diffusion properties of NANOG because we reasoned that abrogating NANOG’s protein-protein interactions should lead to a faster diffusing NANOG. To test this, we repeated SMLFM for NANOG W10A-HALO at 30 ms time resolution, again observing an immobile chromatin-bound population with *D*_app_ of 0.31 ± 0.05 μm^2^ s^−1^ (*α* of 0.73 ± 0.04) and a freely diffusing population with *D*_app_ of 0.98 ± 0.13 μm^2^ s^−1^ (*α* of 0.84 ± 0.05) (**Fig.6B, Supp Fig.3A-D**). As expected, the *D*_app_ of the freely diffusing NANOG W10A mutant was indeed higher than that of wild-type NANOG (p=10^-6^) and similar to that of SOX2, consistent with a loss in NANOG protein complex formation. However, intriguingly, we observed an unexpected reduction in *α* (p=10^-5^), suggesting that the diffusing NANOG W10A molecules were somehow more constrained in their motion within the nucleus. We concluded from this that the W10A mutation was leading to a loss in NANOG’s protein-protein interactions as expected but that it was also somehow constraining the motion of NANOG.

**Figure 6:**
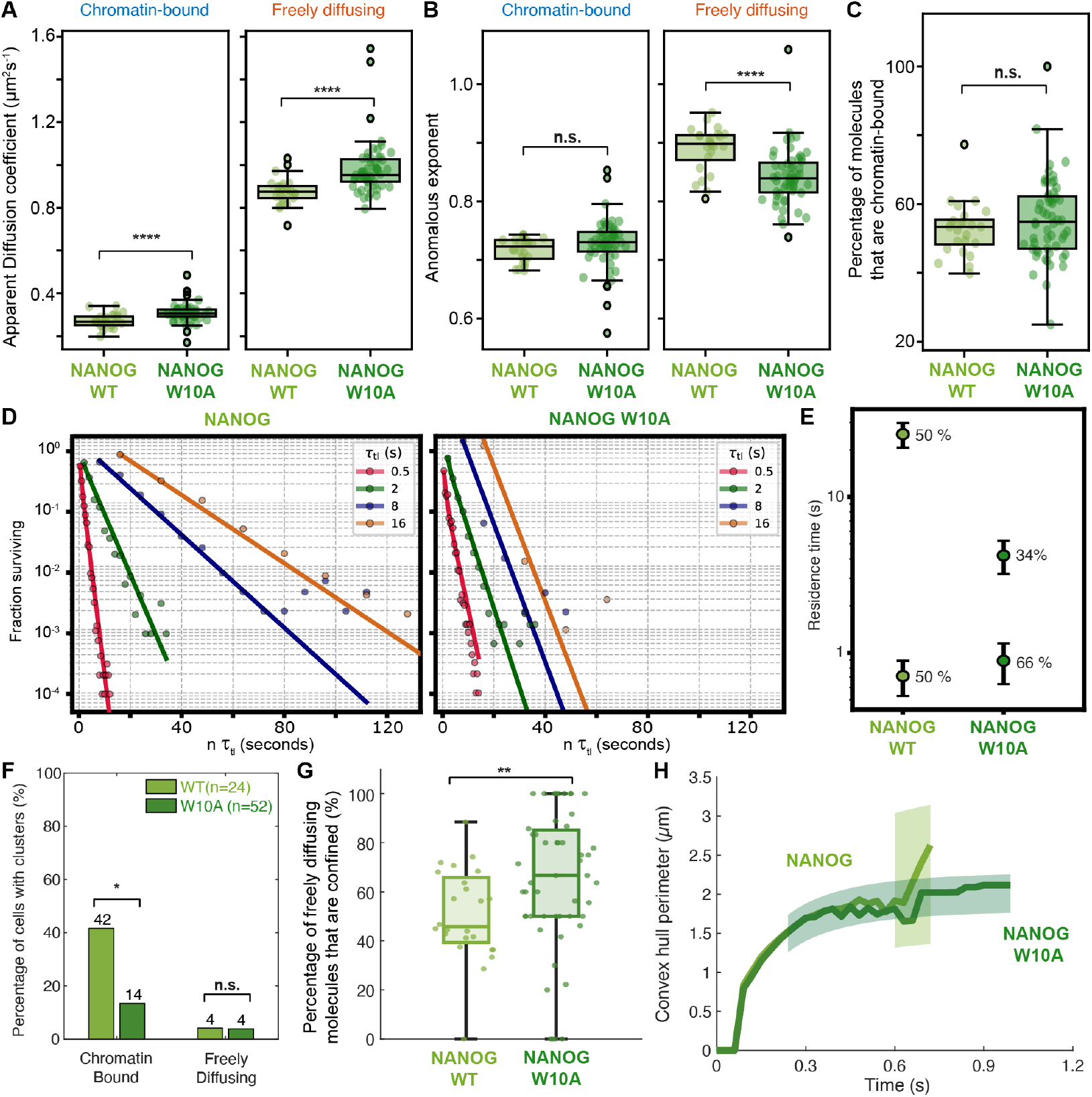
NANOG requires its protein-protein interaction domain to bind DNA robustly and to form larger phase-separated domains. **(A)** Boxplot showing diffusion coefficients of chromatin-bound and freely diffusing wild-type NANOG (NANOG WT) or NANOG with a mutated protein-protein interaction domain (NANOG W10A). Dots represent average values from each fied of view (FOV) (n = 24/56 cells for NANOG WT/W10A). [Mann Whitney, p-values: ****10^-5^ (chromatin bound), ****10^-6^ (freely diffusing)] **(B)** Boxplot showing anomalous exponents of chromatin-bound and freely diffusing NANOG/SOX2. Dots represent average values per FOV. (n = 24/56 cells for NANOG WT/W10A) [Mann Whitney, p-values: 0.09 (chromatin bound), ****10^-5^ (freely diffusing)]. **(C)** Boxplot showing percentage of chromatin-bound NANOG/NANOG W10A (light green/dark green) (n = 24/56 cells for NANOG WT/W10A) [Mann Whitney, p=0.2; Levene’s test for variance p=0.04]. **(D)** Log plot showing fraction of (left) NANOG WT and (right) NANOG W10A molecules surviving at a given time of imaging where the time is the number of frames multiplied by the time lapse between exposures τ_tl_. Datasets for 0.5s, 2s, 8s and 16s time-lapses (blue, orange, green and red) are fit to a single/double exponential with rate constant k_eff_. (for 0.5/2/8/16 s intervals, n = 10/6/7/9 cells for NANOG WT; n = 10/6/4/8 cells for NANOG W10A). **(E)** Dot plots showing chromatin residence times and photobleaching times for NANOG WT and NANOG W10A, calculated by taking the inverse of k_off_ and k_b_, values extracted from global fitting of graphs in (C). **(F)** Barplot comparing number of NANOG WT and NANOG W10A cells exhibiting clustering using the Delaunay-based spatial clustering pipeline (**Fig.4**) [Fisher’s exact test, *p=0.016 for chromatin-bound molecules; p=1 for freely diffusing molecules]. **(G)** Boxplot showing greater percentage of molecules classified as domain-confined at the single-cell level for NANOG W10A (dark green) than for NANOG WT (green) molecules using the convex hull analysis pipeline (n = 24/56 cells for NANOG WT/W10A) [Wilcoxon rank sum test, *p = 0.004]. **(H)** Comparison of the mean and 95% confidence interval of the fitted radii shows a larger and greater range of PSD radii for NANOG WT (green) than for NANOG W10A (dark green). (n = 24/56 cells for NANOG WT/W10A)

To assess how NANOG’s protein-protein interaction domain was influencing chromatin binding, we next examined the chromatin binding kinetics of the NANOG W10A using time-lapse SMLFM. Interestingly, although there was no change in the percentage of chromatin-bound NANOG W10A (56 ± 12%) compared to wild-type NANOG (53 ± 8%), there was now much greater heterogeneity in binding between cells (p=0.04) (**Fig.6C**), suggesting that NANOG’s protein-protein interaction domain was important for ensuring the robustness of NANOG chromatin binding at the single-cell level, possibly explaining why the mutation increases the probability of differentiation (Mullin et al. 2017).

Since NANOG protein complex formation may contribute to this robust binding by increasing its chromatin interaction surface and therefore its residence time, we repeated time-lapse SMLFM for cells expressing the NANOG W10A mutation. This revealed that NANOG W10A datasets fit well to two chromatin binding modes with 66 ± 16% of molecules having a residence time of 0.9 ± 0.3 s and 34 ± 16% of molecules with a residence time of 4.2 ± 1.0 s (**Fig.6D-E, Supp Fig.5**). Although the shorter residence time matched that observed for wild-type NANOG (0.71 ± 0.18 s), the longer one was 6-fold smaller than the value for wild-type NANOG (25 ± 5 s), which was surprising given that we had only mutated NANOG’s protein-protein interaction domain. Assuming that NANOG’s short and long residence times were non-specific and specific binding respectively, we concluded that the W10A mutation had no effect on NANOG’s ability to bind to chromatin non-specifically but that it considerably impaired the time NANOG spent bound once it had found its specific chromatin binding site, likely contributing to the robustness of NANOG binding at the single-cell level. The dramatic decrease in residence time indicated that NANOG’s binding domain contributes little to chromatin binding itself and that its longer 25 s residence time is likely dependent on interactions with itself and/or with other proteins such as SOX2.

### NANOG’s protein-protein interaction domain increases the probability of forming larger PSDs

Since the W10A mutation was altering NANOG’s chromatin binding kinetics and since it was also leading to constraining of diffusing NANOG W10A molecules (lower *α* values), we hypothesised that the W10A mutation was also influencing the formation of PSDs. To examine this, we repeated our clustering analysis. Indeed, this revealed a decrease in the number of cells with clustered chromatin-bound NANOG molecules (p=0.016) but no change in the number of cells with clustered freely diffusing NANOG molecules (**Fig.6F**), suggesting that the primary role of NANOG’s WR domain was to bring DNA binding sites into these PSDs. We also again analysed the freely diffusing NANOG to determine whether the mutation was also changing PSD composition and size. Indeed, this was the case: where previously we had observed 51 ± 19 % of NANOG molecules moving within PSDs, we now found that this had increased to 64 ± 28 % (p=0.004) (**Fig.6G**). Moreover, we found that the radii of these PSDs had now decreased in size from 0.39 ± 0.05 µm to 0.325 ± 0.006 µm (**Fig.6H**), consistent with molecules becoming constrained into smaller PSDs. To test whether this smaller PSD could explain the decrease in *α* value observed above for NANOG W10A trajectories (**Fig.6B**), we simulated trajectories using the average D_app_ and PSD sizes extracted for NANOG and NANOG W10A. Indeed, the increase in D_app_ and decrease in PSD size observed for NANOG W10A was sufficient to lead to a lower *α* value (p=10^-5^) (**Supp Fig.9K**). We concluded from this that NANOG’s tryptophan-repeat domain was increasing the likelihood of forming PSDs and that it was also required for forming larger PSD domains. Interestingly, the large range of radii observed for NANOG encompassed both the larger radii of SOX2-containing PSDs and the smaller radii of NANOG W10A-containing PSDs. This suggested that wild-type cells contained PSDs generated from both NANOG on its own and NANOG in complex with other proteins e.g. SOX2/itself.

**Supplementary Figure 10:**
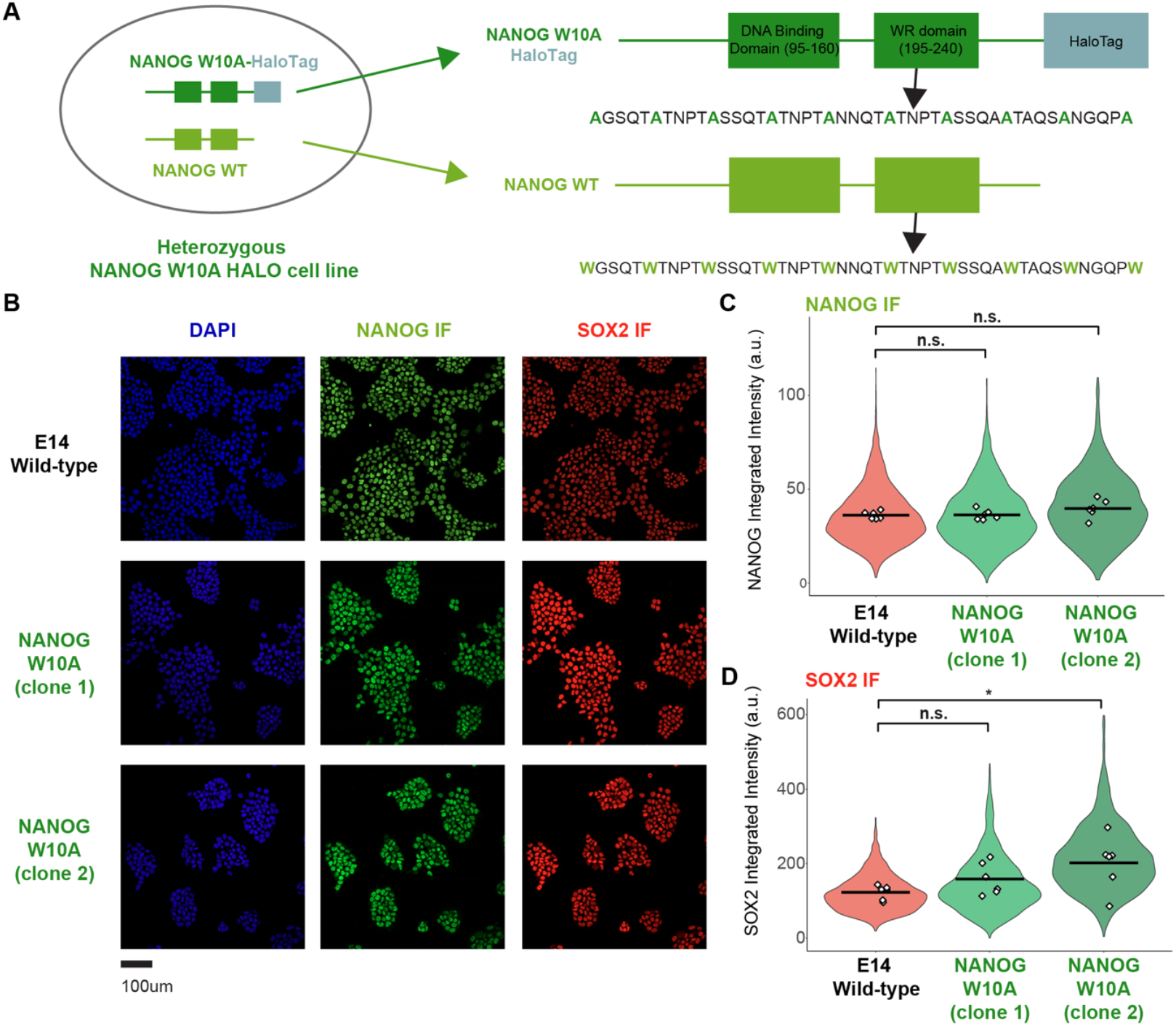
NANOG W10A HaloTag cell line generation. **(A)** Schematic showing heterozygous knock-in cell line where NANOG W10A mutant allele has the fused HaloTag. **(B)** Representative immunofluorescence (IF) images showing DAPI (blue), NANOG (green), and SOX2 (red) staining across three cell lines: E14, W10A clone C1, and W10A clone C2. Scale bar = 100 μm. **(C-D)** Violin plots of single cell measurements showing probability density of **(C)** NANOG and **(D)** SOX2 integrated intensity distribution across cell lines in (A) revealed no significant differences between groups for NANOG expression but a minor increase in SOX2 expression for W10A clone C2. White diamonds represent field-of-view (FOV) means, and black horizontal lines indicate the mean of FOV means. (N=6 FOVs, E14: 4,333 cells, W10A C1: 3,473 cells, W10A C2: 1,841 cells). [one-way ANOVA with Bonferroni-corrected post-hoc pairwise comparisons, for NANOG IF: E14 vs WT10A C1: p = 1, E14 vs WT10A C2: p = 0.3; for SOX2 IF, E14 vs WT10A C1: p = 0.7, E14 vs WT10A C2: *p = 0.04].

## Discussion

In summary, we have used SMLFM to image single TFs for the first time and discovered several advantages including the ability to track TFs throughout the entire nucleus and to collect trajectories rapidly at high density, which allows us to determine the spatial positions of trajectories within a relatively short window of time. Since intra-nuclear PSDs such as pericentromeric heterochromatin and mediator condensates, where NANOG likely binds, are highly dynamic within pluripotent cell nuclei (Du et al. 2024; Novo et al. 2022), this ability to monitor the spatial distribution of TFs rapidly in 3D, and before the PSD has itself moved within the nucleus, provides a considerable advantage. Indeed, using our new approach, we uncovered the spatiotemporal dynamics of the core TF SOX2 and the auxiliary TF NANOG, discovering that they exhibit unique and distinct spatiotemporal strategies.

We discovered that SOX2 and NANOG both diffuse within intra-nuclear PSDs with radii of ∼400 nm, with NANOG using its protein-protein interaction domain to increase the size of its PSDs (**Fig.5-6**). Although clustering of chromatin binding sites has previously been observed (Liu et al. 2014; Okamoto et al. 2023), we show here that diffusing proteins also form part of PSDs (**Fig.5**). The low likelihood of detecting spatial clusters of diffusing molecules (**Fig.4**) may suggest that SOX2/NANOG PSDs are primarily driven by clustering of its chromatin binding sites or that SOX2/NANOG enter PSDs generated by other factors (e.g. mediator) and so they themselves do not cluster despite being restricted in motion within a PSD. Clustering of DNA binding sites is consistent with multiple NANOG binding sites being found at specific genomic regions, e.g. at pericentromeric repeat regions and super-enhancers (Novo et al. 2016). Moreover, single-cell and bulk 3D genomics approaches have previously indicated that NANOG binding sites are spatial clustered with those of several other TFs (e.g. SOX2, TCF3, NR5A2) and that NANOG mediates bridging of its binding sites through 3D chromatin looping through its tryptophan repeat domain (Ma et al. 2018; Novo et al. 2018; Choi et al. 2022; Stevens et al. 2017). Imaging has also previously revealed dispersion of pericentromeric heterochromatin regions in NANOG-null cells (Novo et al. 2016), consistent with a model where NANOG, and specifically its tryptophan-repeat domain, actively contributes to bringing these regions together. How these PSDs are formed requires further study. However, they are consistent with simulations showing that non-specific bridging by TFs lead to clustering of both TFs and their binding sites through an entropic bridging-induced attraction that minimizes entropic penalties arising from the bending and looping of the DNA template (Brackley et al. 2013). NANOG’s tryptophan repeat domain increasing the size of PSDs is consistent with this model.

We also find that, while NANOG is expressed at low levels, it takes advantage of its protein-protein interaction domain to bind DNA for 5-fold longer (**Fig.6**), which compensates for its low expression level. This domain allows NANOG to bind to itself or to core pluripotency TFs such as SOX2 (Mullin et al. 2008; Gagliardi et al. 2013; Mullin et al. 2017) and so this 5-fold increase in NANOG’s residence time may arise from either homotypic self-interactions between NANOG molecules or from heterotypic interactions (e.g. NANOG-SOX2 or NANOG-OCT4). Although simulations reveal that this domain can in principle bring large numbers of NANOG molecules together (Mizutani et al. 2023), heterotypic interactions are likely to be more common (Mistri et al. 2022). Indeed, our observation that NANOG has a much lower concentration than SOX2 (**Fig.1**) would suggest that NANOG’s primary role is to form heterotypic interactions with and stabilise the chromatin residence time of the higher expressing core TFs SOX2/OCT4. This is consistent with NANOG’s role as an auxiliary TF, since it will only add 5 s of additional stability to SOX2/OCT4’s chromatin residence time. It is also consistent with observations that overexpressing NANOG, which increases the likelihood of NANOG oligomerisation, further stabilises naïve pluripotency (Mitsui et al. 2003; Chambers et al. 2003) since this overexpression will drive greater stabilisation of its own chromatin residence time and that of core TFs SOX2/OCT4.

The clustering of NANOG and SOX2 binding sites will considerably influence these chromatin binding kinetics. An oligomerised TF has multiple DNA binding domains and so the probability of losing all protein-DNA interactions prior to a chromatin re-association event is low if there are many other binding sites nearby. This is referred to as a “kinetic trap” where clustering of binding sites increases re-association so that the residence time no longer scales linearly with the number of binding domains within the protein complex. In SMLM measurements, the TF would appear chromatin-bound even though it would actually be alternating between multiple possible combinations of nearby binding sites, effectively hopping from one NANOG binding site to another.

What do these observations mean for the function of NANOG? The clustering of NANOG/SOX2 binding sites is reminiscent of lac repressor where oligomerisation reduces the search time required to rebind at the same binding site (Choi et al. 2008). In the case of lac repressor, this rapid re-association leads to a bistable switch, with rapid reassociation rates driving long residence times and rare losses in chromatin binding leading to long search times. Since NANOG is an activator, its long chromatin residence time and enlargement of PSDs will increase the time genes remain activated in response to external stimuli. NANOG expression levels are likely kept at low levels to limit the likelihood of oligomerisation and to thereby ensure the mix of radii sizes we observe in wild-type cells. Although NANOG overexpression would stabilise PSDs, it would also block the cell’s ability to differentiate in response to signals. Indeed, NANOG-high and NANOG-low cells are known to have different propensities for differentiation (Kalmar et al. 2009; Abranches et al. 2014), possibly because NANOG expression is tightly linked to the probability of forming protein-protein interactions that are essential for the long chromatin residence times and large PSDs.

Notably, while NANOG is dispensable for the maintenance of naïve pluripotency in ESCs, it is required *in vivo* where knockouts result in preimplantation lethality due to a failure to generate naïve pluripotent cells (Silva et al. 2009; Mitsui et al. 2003). One of the key differences is that naïve pluripotent cells emerge transiently *in vivo* while mESCs are grown in self-renewing conditions. *In vivo*, pluripotent cells require NANOG to act rapidly and stably and to outcompete the binding of other TFs such as GATA6 that bind at the same genomic sites (Thompson et al. 2022). In this context, spatial clustering of binding sites and long residence times may block rapid switches in NANOG vs GATA6 binding, stabilising binding modes and thereby cell fate acquisition through a cooperative bistable switch. It is also worth noting that *in vivo*, chromatin is decondensed when NANOG is first expressed and nuclear size dramatically decreases as pluripotent cells emerge so NANOG may play a key role in reorganising chromatin binding sites into PSDs during this phase. Future studies should focus on establishing similar SMLFM experiments *in vivo* to provide deeper insight into this process.

### Beyond pluripotency TFs, we have established unique 3D-SMLM pipelines capable of high-density whole-nucleus spatiotemporal tracking of TFs in a single-cell single-molecule manner

Our pipelines can also be adapted for use in 2D-SMLM and of more than just TFs. We built on our previously published temporal pipelines to now establish and integrate two new spatial pipelines. The first Delaunay-based clustering pipeline separately analysed the spatial clustering of chromatin-bound and freely diffusing TFs, providing insights into how spatial clustering occurs. Moreover, our new convex hull analysis pipeline allowed us to determine how diffusing proteins explore their microenvironment, whether they are part of PSDs and, if so, the size of these PSDs. Since many proteins are expressed at low levels, entering pre-existing PSDs, these pipelines have widespread application. Finally, our analysis pipelines output spatial and temporal metrics for TFs at the single cell level, which may prove useful when there is greater cell heterogeneity, for example during asynchronous stem cell differentiation processes.

## Methods

### Plasmids used

Plasmids used:
| Plasmid Name | Description | Source |
| --- | --- | --- |
| NANOG-HALO targeting vector | Knock-in targeting construct comprising 800 bp homology arms flanking a HaloTag-TEV-Avi-FLAG sequence and blasticidin resistance cassette (surrounded by Rox sites) to enable stable expression of NANOG-Halo-TEV-Avi-FLAG fusion protein. The HaloTag sequence is targeted to the 3' end of the gene locus replacing the endogenous STOP codon. | (this study) |
| SOX2-HALO targeting vector | Knock-in targeting construct similar to NANOG-HALO vector but to enable stable expression of SOX2-Halo fusion protein. | (this study) |
| NANOG(W10A)-HALO targeting vector | Knock-in targeting construct similar to NANOG/SOX2-HALO vector but incorporating W10A mutations into homology arm to block NANOG oligomerisation. Vector does not include blasticidin resistance cassette. | (this study) |
| Dre recombinase | Plasmid expressing Dre recombinase to remove blasticidin cassette. | (Basu et al. 2023) |

### Mouse embryonic stem cell (mESC) lines

Mouse embryonic stem cell (mESC) lines:
| Name in manuscript | Description | Source |
| --- | --- | --- |
| ESCs | ES-E14tg2A mESC line derived from a male mouse strain 12910la. | (Ying et al. 2008) |
| *NANOG-HALO* | ES-E14tg2A mESC line with knock-in of HaloTag to C-terminus of *Nanog* gene locus. | (this study) |
| *NANOG (W10A)-HALO* | ES-E14tg2A mESC line with knock-in of HaloTag to C-terminus of *Nanog* gene locus. | (this study) |
| *SOX2-HALO* | mESC line with knock-in of HaloTag to C-terminus of Sox2 gene locus. Knock-in carried out in F1G4 background mESC line expressing two reporter constructs T<sup>mCherry</sup> and Sox2<sup>Venus</sup> (Koch et al. 2017) | (this study) |

### Software

#### Data collection

Microscope image acquisition

Micro-manager (Edelstein et al. 2014)

(https://www.micro-manager.org)

ImageJ software

(Schneider, Rasband, and Eliceiri 2012)

LAS X (Leica Application Suite X) (Leica Stellaris software)

#### Data analysis - imaging

CellProfiler (Stirling et al. 2021)

https://cellprofiler.org/

SMLFM peak fitting (Daly et al. 2024)

https://github.com/Photometrics/PySMLFM

Trajectory analysis (Basu et al. 2018)

https://github.com/wb104/trajectory-analysis

Trajectory classification (Basu et al. 2023)

https://zenodo.org/records/17056157

Data plotting and stats R

R Core Team (2023).

R: A language and environment for statistical computing. R Foundation for Statistical Computing, Vienna, Austria. URL https://www.R-project.org/.

### ESC culture conditions

ESCs were grown in phenol red-free 2iLIF medium at 37 °C, 7% CO_2_ and 80% humidity as previously described (Ying et al. 2008). Phenol red-free 2iLIF medium was prepared fresh every week by adding 1 µM PD0325901 (ABCR, AB 253775), 3 µM CHIR99021 (ABCR, AB 253776) and 10 ng/ml murine leukaemia inhibitory factor (mLIF, provided by the Department of Biochemistry, University of Cambridge) to N2B27 basal medium. N2B27 medium is composed of 50% phenol red-free DMEM/F-12 (Gibco #21041025) and 50% phenol red-free and L-Glutamine-free Neurobasal (Gibco #12348017) supplemented with 0.5x N2 (made and batch tested in-house by Cambridge Stem Cell Institute), 1x B-27− Supplement (Gibco #17504044), 2mM L-Glutamine (Life tech, #25030024), and 0.1 mM 2-mercaptoethanol (in-house/Life Tech, #1985023). Cells were passaged every two days by washing in PBS (Sigma-Aldrich #D8537), followed by incubation with Accutase (Biolegend, 423201) for 2 min at 37 C to detach. Cell clumps in suspension were then washed and pelleted in PBS before resuspending and re-plating as single-cell suspension in fresh medium. To help cells attach to the surface, plates were incubated for 15 minutes at 37 C in PBS containing 0.2% gelatin (Sigma-Aldrich #G1890). All cell lines were routinely screened for mycoplasma contamination at least twice yearly and tested negative.

### Cell line generation

ESCs expressing NANOG/SOX2 tagged at the C-terminus with HaloTag were generated by CRISPR-Cas9 knock-in of a cassette containing HaloTag-Flag and a blasticidin selection gene into one allele. The blasticidin cassette, which was flanked by Rox sites, was then removed using Dre recombinase. ESCs expressing NANOG W10A tagged at the C-terminus with HaloTag were generated by CRISPR-Cas9 knock-in of a cassette containing HaloTag-Flag followed by labelling with JFX_554_ dye (see below) and fluorescence-activated cell sorting of single positive clones into a 96-well plate. Since knockout of *NANOG/SOX2* is lethal, we used cell viability assays, RT-qPCR of NANOG/SOX2-dependent genes and quantitative immunofluorescence to verify that the function of the tagged NANOG/SOX2 was not impaired. Western blots were carried out to confirm heterozygous vs homozygous expression.

### RT-qPCR

A cell pellet was collected from a 6-well plate in 400 μl of TRIzol (Invitrogen, 15596026) and RNA extracted using the Direct-zol RNA MiniPrep Plus kit (Zymo Research, R2072). A NanoDrop One spectrophotometer (Thermo Fisher Scientific, ND-ONE-W) was used to measure RNA concentration. Reverse transcription was performed on 500ng of RNA using the SuperScript IV First-Strand Synthesis kit (Thermo Fisher Scientific, 18091050). 25 ng of cDNA was used for qPCR alongside 0.5 μl of the VIC-labelled Gapdh control probe (Thermo Fisher Scientific, 4352339E) and 0.5 μl of the following tareget-specific FAM-labelled Taqman probes (Thermo Fisher Scientific, 4448892):

| Taqman Gene Expression Assay | Assay ID |
| --- | --- |
| *Esrrb* | Mm00442411_m1 |
| *Nanog* | Mm02384862_g1 |
| *Otx2* | Mm00446859_m1 |
| *Sox2* | Mm03053810_m1 |

The qPCR reaction was performed in triplicate using the Taqman Universal PCR Master Mix (Thermo Fisher Scientific, 4366072) and loaded onto a StepOnePlus Real-Time PCR System (Thermo Fisher Scientific, 4376600). Raw data was exported as a .csv file and analysed in Microsoft Excel following the ΔΔCt method.

### Immunofluorescence

Cells were grown on 8-well micro-chamber slides (Ibidi, IB-80826) and fixed for 10 min using 4% para-formaldehyde (PFA; Thermo Fisher Scientific, 28906) at room temperature (RT). Cells were washed twice with PBS and then blocked and permeabilised for 2 hours in blocking buffer (0.1% Triton-X (Sigma-Aldrich, BCCB0860) + 3% donkey serum (Sigma-Aldrich, D9663) in PBS). Samples were incubated at 4°C overnight with primary antibodies diluted in blocking buffer followed by three washes in PBST (PBS + 0.1% Triton 100X). Samples were then incubated with secondary antibodies and 1 µg/ml of DAPI (Sigma-Aldrich, MBD0015) diluted in blocking buffer for 40 min in the dark and at RT. This was followed by three washes in PBST and a final wash into PBS for storage at 4°C prior to imaging. A Nikon Ti inverted microscope with a Yokogawa spinning disk scanhead and a Photometrics BSI Prime camera fitted was used to collect 3 x 3 tile scan micrographs using a 20x objective lens.

### List of primary antibodies used in this study

List of primary antibodies used in this study:
| Antigen | Species | Concentration | Manufacturer | Cat # | LOT |
| --- | --- | --- | --- | --- | --- |
| NANOG | rabbit | 1:200 | Abcam | ab80892 | A300397A2 |
| SOX2 | rat | 1:200 | Invitrogen | 14981182 | 2493179 |
| NANOG | rabbit | 1:200 | Bethyl Laboratories | A300-397A | A300-397A-2 |
| ESSRB | mouse | 1:300 | Perseus | PP-H6705-00 | A-2 |

### List of secondary antibodies used in this study

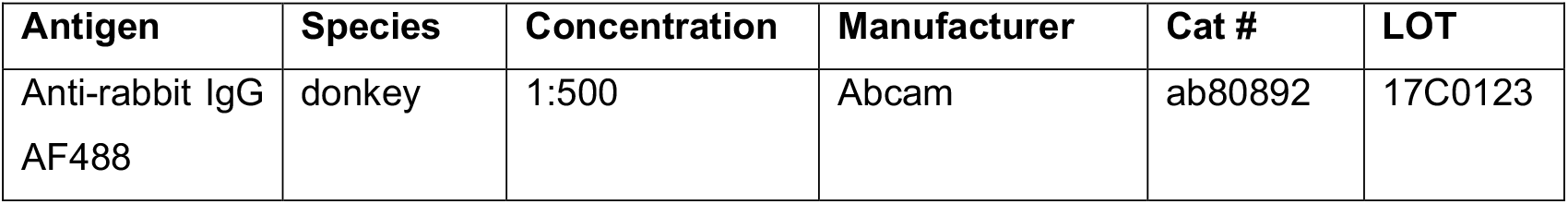
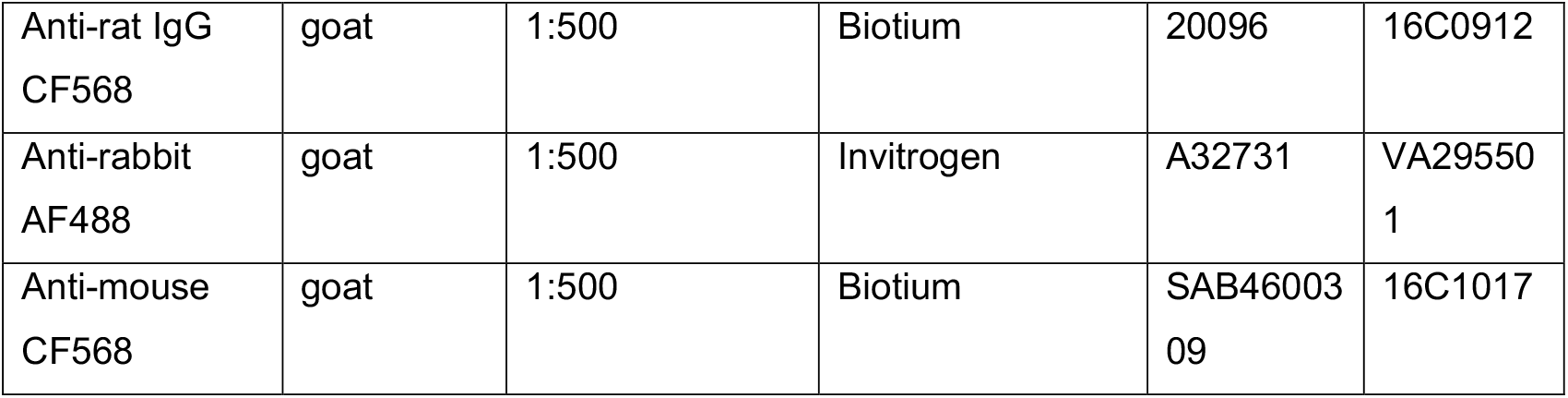

List of secondary antibodies used in this study:
| Antigen | Species | Concentration | Manufacturer | Cat # | LOT |
| --- | --- | --- | --- | --- | --- |
| Anti-rabbit IgG AF488 | donkey | 1:500 | Abcam | ab80892 | 17C0123 |
| Anti-rat IgG<br>CF568 | goat | 1:500 | Biotium | 20096 | 16C0912 |
| Anti-rabbit<br>AF488 | goat | 1:500 | Invitrogen | A32731 | VA29550<br>1 |
| Anti-mouse<br>CF568 | goat | 1:500 | Biotium | SAB46003<br>09 | 16C1017 |

### Microscope Settings for spinning disk

Microscope Settings for spinning disk:
| Fluorophore | Excitation<br>Wavelength | Laser<br>Power<br>(%) | Exposure<br>Time |
| --- | --- | --- | --- |
| DAPI | 405 nm | 10 | 1 s |
| AF488 (NANOG) | 488 nm | 20 | 1 s |
| AF568 (SOX2) | 561 nm | 5 | 1 s |

For validation of NANOG W10A HALO ESCs, confocal microscopy was performed using a Leica STELLARIS 5 (DMI8-CS) microscope with LAS X software version 4.8.2.29567 and an HC PL APO CS2 20x/0.75 numerical aperture dry objective. Images were acquired in 1024 x 1024 pixel format with 0.57 μm pixel size, resulting in a 581.25 μm x 581.25 μm field of view. Acquisition parameters included unidirectional scanning at 400 Hz with 1.4 μs pixel dwell time, and 16-bit photon counting detection mode using HyD S silicon photomultiplier detectors.

### Microscope Settings for Stellaris Confocal

Microscope Settings for Stellaris Confocal:
| Target | Fluorophore | Excitation<br>(nm) | Laser<br>Power<br>(%) | Emission<br>Range<br>(nm) | Detector | Gain | Line<br>Averaging |
| --- | --- | --- | --- | --- | --- | --- | --- |
| DAPI | DAPI | 405 | 7 | 410-460 | HyD S 1 | 25 | 2 |
| NANOG | AF488 | 499 | 18 | 520-540 | HyD S 2 | 33 | 3 |
| SOX2 | AF568 | 555 | 19 | 580-611 | HyD S 3 | 15 | 2 |
| ESRRB | CF568 | 555 | 20 | 580-611 | HyD S 3 | 30 | 2 |

### Quantification of NANOG/SOX2-Halo protein levels

For quantification of NANOG/SOX2 protein levels, we carried out live-cell imaging of Halo-JFX_549_-labelled samples. Poly-D-Lysine coated 35 mm glass bottom (#1.5) dishes (MatTek Corporation, #P35G-1.5-14-C) were coated with 200 µl of fibronectin (20 μg/ml, Sigma-Aldrich, #F0895) for 30 minutes. The fibronectin solution was then aspirated, and the dishes left in the incubator for 30 minutes to air dry. 50,000 naïve ESCs were seeded (seeding density of 5,191 cells/cm^2^) on dried MatTek dishes and cultured in 2 ml of 2iLIF media for 48 hours prior to imaging. Cells were labelled by adding 500 nM Halo ligand dyes (JFX_549_) the evening before imaging and incubating cells in dye overnight in the incubator (dyes from Janelia Research Campus; reconstituted in DMSO) (Grimm et al. 2016; Grimm et al. 2015). For nuclear labelling and segmentation, cells were labelled with 20 nM of Hoechst dye at the same time as adding the Halo dye. Just before imaging, unbound ligands were washed as previously described (Steindel et al. 2022): cells were first washed in PBS (Sigma-Aldrich, #D8537) and then cultured for 30min in fresh 2iLIF media. The media was then changed again prior to imaging. A Nikon Ti inverted microscope with a Yokogawa spinning disk scanhead and a Photometrics BSI Prime camera fitted was used to collect 2-channel (JFX_549_ and Hoechst) 3 x 3 tile scan micrographs using a 20x objective lens.

### Microscope Settings for spinning disk

Microscope Settings for spinning disk:
| Fluorophore | Excitation Wavelength | Laser Power | Exposure Time |
| --- | --- | --- | --- |
| Hoechst | 405 nm | 10.0% | 1 s |
| JFX<sub>549</sub> | 561 nm | 15.0% | 1 s |

### Live-cell 3D Single Molecule Light-Field Microscopy (SMLFM)

ESCs were cultured on fibronectin-coated 35 mm glass-bottom dishes (MatTek Corporation, P35G-1.0-14-C) as described previously. Cells were labelled with the HaloTag-PA-JF_646_ dye 12–16 h before imaging to allow sufficient binding time (500 nM for 30 ms exposure experiments and 50 nM for 500 ms experiments). Unbound dye molecules were removed by two PBS washes, followed by incubation in imaging medium at 37 °C for 30 minutes, and imaging was performed in fresh media.

To extract chromatin binding kinetics and spatial clustering metrics, SMLFM was then conducted at 30 ms exposure as previously described (Daly et al. 2024). Briefly, it incorporated an epi-fluorescence microscope (Eclipse Ti-U, Nikon) equipped with a 1.27 NA water immersion objective lens (Plan Apo VC 60×, Nikon), with z-position control via a scanning piezo (P-726 PIFOC, PI). The setup included a 4f lens configuration, a hexagonal microlens array (f = 175 mm, pitch = 2.39 mm, custom-made by CAIRN Research), and an EMCCD camera (Evolve Delta 512, Photometrics). Excitation was achieved using 638 nm (75 Wcm^-2^) and 405 nm lasers (0.003 Wcm^-2^), both circularly polarised, collimated, and focused onto the back focal plane of the objective. Fluorescence was collected by the same objective and separated from the excitation beam using a quad-band dichroic mirror (Di01-R405/488/561/635-25 ×36, Semrock). Emission filters (BLP01-647R-25, Semrock; FF02-675/67-25, Semrock) were placed immediately before the detector to isolate fluorescence emission. Laser powers were 450 Wcm^-2^ and 0.003 Wcm^-2^ for the 638 nm and 405 nm lasers respectively.

For residence time experiments, we implemented a time-lapse imaging approach that we have previously established to extract the residence time of a TF on chromatin while accounting for photobleaching (Gebhardt et al. 2013; Basu et al. 2023). SMLFM was performed with a fixed exposure time of 200 ms (to motion blur freely diffusing molecules) interspersed with dark intervals of varying lengths (0.3, 1.8, 7.8, and 15.8 s). SMLFM was conducted on a separate SMLFM microscope that was set up as above but with different 638 nm (180 mW, 06-MLD 638 nm, Cobolt) and 405 nm lasers (365 mW, 06-MLD 405nm, Cobolt). We also modified the detection path with a different hexagonal microlens array (f = 100 mm, pitch = 2.39 mm, PowerPhotonic Ltd) and a Kinetix sCMOS camera (1T-01-KINETIX-M-C, Photometrics). Laser powers were 12.5 Wcm^-2^ and 0.003 Wcm^-2^ for the 638 nm and 405 nm lasers respectively.

### Quantification and statistical analysis-Extracting NANOG/SOX2 levels from HALO-JFX_549_ labelling and quantitative immunofluorescence datasets

Images were visualised using ImageJ (Schneider, Rasband, and Eliceiri 2012) and analysed using CellProfiler (Stirling et al. 2021). CellProfiler was used to segment nuclei from the Hoechst/DAPI channel and then to measure the signal intensity of the other channels. This intensity was normalised using the Hoechst/DAPI signal to account for differences in cell size or cell cycle state. Here is a table of the optimised Cell Profiler pipeline for NANOG and SOX2 protein quantification from Halo-JFX_549_ labelling, showing the six modules used, in order, and the settings in each module that were altered from the default:

| Module | Custom Setting | Value |
| --- | --- | --- |
| CorrectIlluminationCalculate | Select how the illumination function is calculated | Background |
|  | Block size | 100 |
|  | Rescale the illumination function? | No |
|  | Smoothing method | Smooth to Average |
| CorrectIlluminationApply | Select how the illumination function is applied | Subtract |
| RescaleIntensity | Rescaling method | Choose specific values to be reset to the full intensity range |
|  | Intensity range for the input image | 0.000 – 0.035 |
| IdentifyPrimaryObjects | Use advanced settings? | Yes |
|  | Typical diameter of objects in pixel units | 30 - 75 |
|  | Threshold strategy | Adaptive |
|  | Thresholding method | Otsu |
|  | Lower and upper bounds on threshold | 0.0005 – 1.0 |
|  | Size of adaptive window | 100 |
|  | Log transform before thresholding? | Yes |
|  | Method to draw dividing lines between clumped objects | Propagate |
|  | Speed up by using lower-resolution image to find local maxima? | No |
|  | Fill holes in identified objects? | After declumping only |
| MeasureObjectIntensity | None | N/A |
| MeasureObjectSizeShape | Calculate the Zernike features? | No |

For analysis of quantitative immunofluorescence datasets, the pipeline was modified slightly:

| Module | Setting | Value |
| --- | --- | --- |
| RescaleIntensity | Intensity range for the input image | 0.00025 – 0.00150 |
| IdentifyPrimaryObjects | Fill holes in identified objects? | After both thresholding and declumping |

Image analysis was performed in *CellProfiler* (v5). The DAPI channel was smoothed with a Gaussian filter (artifact diameter set to 2 pixels) and used to segment nuclei with the IdentifyPrimaryObjects module (typical nuclear diameter: 15–32 pixels). Segmentation was refined using adaptive Otsu thresholding with shape-based declumping, excluding border-touching and out-of-range objects. The resulting nuclear masks were applied to quantify single-cell fluorescence intensity of NANOG, SOX2, ESRRB, and H3K9me3 using the MeasureObjectIntensity module. Measurements were exported as a .csv file for downstream statistical analysis.

### SMLFM image analysis

A Jupyter Notebook-based pipeline was developed, integrating Python and MATLAB scripts for automated processing and analysis of SMLFM data. A detailed guide to implement this can be found here: https://github.com/diyanj03/SMLFM_Analysis_DJ

Briefly, the pipeline does the following: (1) Fits 2D localisations, (2) Calculates 3D localisations from the multiple perspective views, (3) curates 3D localisation datasets, (4) tracks 3D localisations over time, (5) extracts biophysical parameters, (6) runs Delauanay-based spatial clustering analysis. Further details about each step is as follows:

#### Step 1: 2D fitting of single molecule images

Raw single-molecule imaging data, acquired as .tif stacks, were initially processed using our ‘fitting_2D.py’ module which relies on the ‘PyImageJ’ python package to interface with Fiji. Processing began by subtracting the average intensity Z-projection to each frame to remove background, followed by addition of a constant offset (value=400 for 30ms and value=100 for 500 ms datasets) to all pixels. Fiji’s ‘Gaussian Blur’ filter (sigma = 0.4) was then applied across the stack to smooth out noise. 2D localisations were then obtained using Fiji’s GDSC SMLM 2.0 PeakFit function (Etheridge, Carr, and Herbert 2022). We used the least-squares solver (fit_solver = LVM LSE) to fit a Gaussian PSF model (psf_model = Circular Gaussian 2D), outputting 2D localisations with intensity and uncertainty estimates across all frames. For SMLFM datasets, this fits the 2D localisations in all perspective views. Parameters for PeakFit were configured as follows:

*Calibration: 266 nm for 30ms and, because of 2×2 binning, 432 nm for time-lapse datasets; Exposure Time: 30 ms or 200 ms; PSF Model: Circular Gaussian 2D; Spot Filter Type: Difference; Spot Filter: Mean; Smoothing: 0*.*80; Spot Filter 2: Mean; Smoothing 2: 2*.*00; Search Width: 1; Border Width: 1; Fitting Width: 4; Fit Solver: LVM LSE; Fail Limit: 3; Pass Rate: 0*.*50; Neighbour Height: 0*.*30; Residuals Threshold: 1; Duplicate Distance: 0*.*50; Shift Factor: 1; Signal Strength: 3 for 30ms and 1*.*5 for time-lapse datasets; Min Photons: 1; Min Width Factor: 0*.*50; Max Width Factor: 2; Precision: 120; Camera Bias: 400 ADU for 30ms and 100 ADU for time-lapse datasets; Gain: 40 electrons/ADU for 30ms and 6 electrons/ADU for time-lapse datasets; Read Noise: 7*.*1 ADU for 30ms and 1*.*2 ADU for time-lapse datasets; PSF Parameter 1: 1*.*700; Relative Threshold: 1*.*0E-6; Absolute Threshold: 1*.*0E-16; Parameter Relative Threshold: 0*.*0; Parameter Absolute Threshold: 0*.*0; Max Iterations: 20; Lambda: 10*.*0000; Precision Method: Mortensen; Image Scale: 1; Image Size: 512; Image Pixel Size: 5*.

#### Step 2: 3D reconstruction of SMLFM 2D data

Our pipeline then carried out 3D reconstruction of SMLFM data using the PySMLFM package (https://github.com/Photometrics/PySMLFM), as previously described (Daly et al. 2024). Briefly, this package first identifies groups of 2D localisations from different perspective views that most likely originate from a single molecule. If a group contains at least four such 2D localisations, the emitter’s 3D position is then calculated by finding the best fit (least-squares estimate) to an optical model that relates the emitter’s axial depth to the observed parallax between these views. A 3D fit was accepted if its residual error was below 200 nm; the contributing 2D localisations were then removed, and this iterative procedure of grouping, 3D fitting, and candidate exclusion was repeated to find and localise subsequent single molecules.

#### Step 3: Curation of 3D localisation datasets

Next, we used the ‘processing_3Dlocs.py’ module from our pipeline to curate each 3D localisations dataset based on specific criteria: (1) localisations on the coverslip, if present, were cropped out by excluding all data below a defined z-coordinate, (2) localisations external to the nuclear boundaries were excluded and cropped out, (3) datasets exhibiting very low localisation density (<5000 localisations in 10,000 frames) or a spatial distribution inconsistent with nuclear morphology (e.g. mitotic cells) were entirely omitted from subsequent analysis.

#### Step 4: Tracking of 3D localisations over time

Using processed 3D localisations, the trajectories of individual molecules were assembled using a custom Python code for connecting localisations in subsequent frames if they were within 800 nm (https://github.com/wb104/trajectory-analysis). This code also outputs average signal intensity per trajectory and trajectory lengths (OPTION - savePositionsFramesIntensities).

#### Step 5A: Classification of trajectories into two diffusion modes

To classify trajectories, we use ‘analyze-tracking-data’, a pipeline we recently established that uses a sliding window along each trajectory to output four biophysical parameters for each sub-trajectory – anomalous exponent (α), apparent diffusion coefficient (D_app_), length of confinement (L_c_) and drift magnitude (https://zenodo.org/records/17056157) (Basu et al. 2023). The algorithm then employs a Gaussian mixture model to classify and segment single molecule trajectories, specifically those exhibiting switching behaviour between different diffusion states, into domain-confined and unconfined states that correspond to chromatin-bound and freely diffusing molecules at 30 ms exposure times. Analysis was carried out using a minimum trajectory length of 5 frames (minNumPoints = 5), a sliding window of 11 frames (numMSDpoints = 5) and 0.03 s exposure time.

#### Step 5B: Residence time analysis

Residence time analyses were conducted using the ‘main_sequentialFit’ and ‘main_globalFit’ modules of the custom ‘residenceTimeAnalysis.py’ script. Briefly, the code extracted trajectory lengths from the -savePositionsFramesIntensities output files. Single-, double- and triple-exponential decay fits (Eq.3, Eq.4 and Eq.5) were then applied to assess whether the data were better described by one, two or three binding populations:

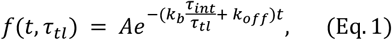

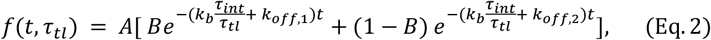

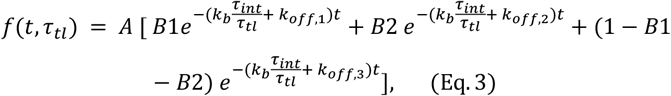

In the double exponential fit, B denotes the fraction of molecules dissociating with rate, k_off,1_. In the triple exponential fit, B1 denotes the fraction of molecules dissociating with rate, k_off,1_ and B2 denotes the fraction of molecules dissociating with rate, k_off,2_. Default parameters were used with ‘min_residence_time’ set to 0.5 to match the shortest time-lapse interval dataset. The fit with lowest Bayesian Information Criterion (BIC) value was used to determine the number of exponentials, *N*, that best fit the data.

#### Step 6: Delaunay-based clustering of live-cell trajectories

To classify trajectories into spatially clustered or non-clustered populations, we employed the ‘Delaunay.py’ module, an approach based on Delaunay triangulation and Gaussian mixture modelling (GMM). This module utilizes the “clusterLightFieldData” MATLAB script to perform the clustering analysis on the motion classification outputs from **Step 5A**. It calls the “clusteringByTriangulationGmm” class to perform Delaunay triangulation on trajectory average positions in single cells and extract the log_10_-normalised median Delaunay edge lengths as features for Gaussian mixture modelling. It first collects an ensemble of m trajectories *X*_*i*_(1), …, *X*_*i*_(*n*_*i*_Δ*t*), where *n*_*i*_ is the number of frames for *i* trajectory (*i* = 1, …,m). To represent the average spatial localizations of these trajectories, it computes the centre of mass (CoM) of each trajectory:

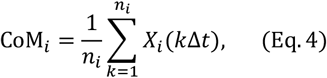

The centre of mass for all trajectories was collected as a set of points CoM = {CoM_1_, …, CoM_m_} and this set of points was used to construct the Delaunay Triangulation D(CoM). As a result, for each trajectory *i* represented by the point CoM_*i*_, it obtains a collection of T_*i*_ triangulation edges 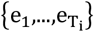 attached to the point. It computes the Euclidean lengths of these edges:

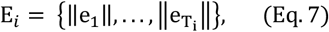

and then gets the 50% percentile (median) edge length ℳ_*i*_ = median(E_*i*_). Finally, it takes the log_10_-normalised median edge length, log(ℳ_*i*_), and collects this feature from all trajectories as a feature vector R to perform GMM:

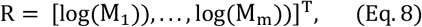

where m is the number of trajectories, and T is taking the transpose. It next performs GMM to separate the feature vector R into several Gaussian distributions. To determine the optimal number of Gaussian, it imposes GMM to separate R into 1 to 10 populations respectively, and for each calculates the associated Bayesian information criterion (BIC) to determine the optimal number of Gaussian components. The minimum BIC gives the best GMM with the optimal number of populations. In the case of monotonously decreasing BICs with increasing number of populations, we selected the GMM with the greatest decrease in the BIC; in other cases where a true minimal BIC existed, we selected the GMM with the lowest BIC. Eventually, we obtained a grouping of trajectories based on their median edge lengths attached in the Delaunay graph D (CoM).

To address the spatial density variation among single cells, we first collected clustering analysis outputs from all cells using “CollectAnalysisMatObjects”. Then, we contextualised the Gaussian components in each cell by using “gmmAnalysis_NanogSox2(“mean”, 11)” to compute a global Gaussian mixture model on the means of the Gaussian components from all cells. The Gaussian components of each cell were reassigned based on the global Gaussian mixture model. Those components assigned to the Gaussian with the smallest mean (i.e. with the most density) in the global Gaussian mixture model were considered spatially clustered, and the other trajectories were assigned as non-clustered.

### Classifying trajectories into two exploration states based on convex hull statistics

To identify trajectories confined by nuclear domains, we developed a new pipeline based on convex hull statistics (https://zenodo.org/records/17036360). The datasets of diffusing molecules were analysed by using “AnalyzeTrajConvexHull”, which computes the time-dependent convex hull perimeter curves of trajectories, performs Dynamic Time Warping (DTW)-K-mean clustering on these curves to distinguish trajectories as domain-confined or unconfined, and finally estimates the radius of the domain by fitting to the appropriate equation (trajectories were projected into the x-y plane to fit this equation). We used a time step of 0.03 s (dt = 0.03) and kept long trajectories with more than 10 frames for analysis (minframe = 10). We applied a variation threshold of 20% (threshold = 0.2) to identify the plateau region of the mean perimeter curve for fitting the asymptotic formula. The steps in detail are as follows:

#### Step 1: Computing the convex hull time course of trajectories

We start with an ensemble of *m* trajectories with position *X*_*i*_(0),…,*X*_*i*_(*n*_*i*_Δ*t*), where *n*_*i*_ is the number of frames for *i* trajectory (*i* = 1,…,*m*) acquired at a time step Δ*t* that we wish to classify into domain-confined and unconfined. For each trajectory *i*, we generate the time-evolving convex hull up to time *t*_*k*_: **CH**_*i*_(*t*_*k*_) = conv(*X*_*i*_(0),..*X*_*i*_(*k*Δ*t*)), from which we compute the perimeter *L*_*i*_(*t*_*k*_) as the sum of external edges:

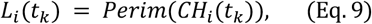

The time-dependent perimeter curves *L*_*i*_ = (*L*_*i*_(0),…,*L*_*i*_(*t*_*ni*_)) for the ensemble L of all *m* trajectories indicate possible confined behaviours

#### Step 2: Classifying trajectories into domain-confined and unconfined populations

To classify trajectories into two classes domain-confined (C) and unconfined (U) populations, we grouped the curves *L*_*i*_ into two clusters. To obtain such clustering, we used the framework of a traditional k-means algorithm (Pelleg and Moore 1999; Lloyd 1982) by adding the following adjustment: we changed the distance metric from Euclidean distance to Dynamic Time Warping (DTW), which accounted for the time-shifts and variant lengths of the curves. We employed this DTW-K-mean algorithm (implemented in Python *<u>tslearn</u>* package) in our convex-hull pipeline. It works by initially seeding the centroids *µ*_1_ and *µ*_2_ of the two clusters (Arthur and Vassilvitskii 2007) by choosing two curves randomly from the ensemble L. The algorithm proceeded by alternating between the two steps:

1. **Assigning step:** assign each perimeter curve to the cluster with the nearest centroid. We use the DTW distance to measure the distance between each perimeter curve *L*_*i*_ to the centroid curve *µ*_*s*_ (*s* = 1,2) (Sakoe and Chiba 1978). The distance is given by:

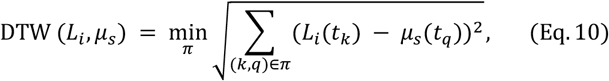

where *t*_*k*_ is the time point associated with the k frame and the alignment of the two curves is encoded in the array

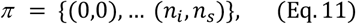

for *n*_*i*_ points in trajectory i, and *n*_*s*_ is the number of points in the centroid. Then, the perimeter curve is assigned to the cluster with a closer centroid:

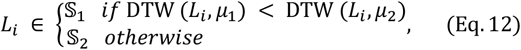
2. **Updating step:** Here the centroid was recomputed for the perimeter curves assigned to each cluster as follows: the centroid *µ*_*s*_ is calculated by the DTW barycenter averaging (DBA) algorithm (Petitjean, Ketterlin, and Gançarski 2011), which minimizes the sum of squared DTW distances between the perimeter curves assigned to the cluster to the centroid:

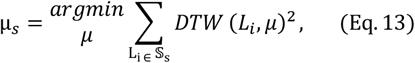

Once the algorithm converged to the final centroids 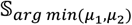, we assigned the trajectories in the smaller cluster as domain-confined (𝒞), and the other trajectories as unconfined (𝒰):

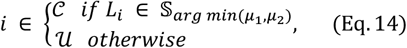

After Step 2 classification, trajectories in cluster 𝒞 had, on average, lower perimeter curves and more confined behaviour than those in cluster 𝒰.

#### Step 3: Extracting the mean radius of domain-confined trajectories

After each trajectory had been classified, we recovered their spatial position and estimated the radius 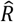 of the confined disc by fitting the mean behaviour computed as:

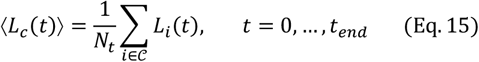

where 𝒞 is the ensemble of indices associated with the domain-confined trajectories, *N*_*t*_ is the number of trajectories that have survived to time *t* until the end time point *t*_end_ of the longest trajectory in the ensemble of domain-confined trajectories.

To extract the radius of confinement *R*, we then used the asymptotic formula for the mean convex hull perimeter *f*(*t,R,D*) for large time *t*. This formula has been derived in the context of Brownian trajectories with diffusion coefficient *D* confined in a disc (De Bruyne et al. 2022). It is given by:

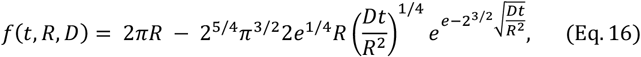

To define the time point *t*_fit_ to start the fit, we identified the longest possible segment which did not vary from its mean LÄ beyond a threshold *β*:

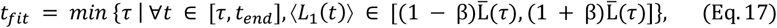

where 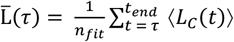, and *n*_fit_ is the number of time points in the segment.

To estimate the radius of confinement 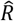, we separately estimated the diffusion coefficient 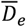 and then used the least-square fitting method on (Eq.16):

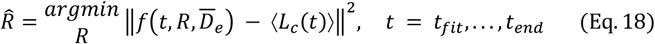

where ∥·∥ is the *L*^2^-norm.

We note that the mean 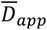 can be computed by averaging the mean squared displacement over the ensemble of trajectories:

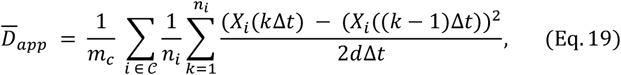

where *m*_C_ is the number of trajectories in C, *n*_*i*_ is the number of frames in trajectory *i*, and *d* is the dimension of the space.

### Simulations to test convex hull method

To validate and evaluate the algorithm, we used “AggregateDiffusionSimulator” to generate synthetic Brownian trajectories inside and outside of a reflective domain and used this ensemble of trajectories with a known disc radius as the ground-truth to evaluate the accuracy of our convex-hull pipeline. Specifically, we simulated a nucleus of 10 µm in radius (radiusOut = 10) containing one reflective domain (numDomains = 1) of 1 µm in radius (radiusIn = 1). We set 100 diffusing particles both inside and outside the domain (numParticleIn = 100; numParticleOut = 100) with random initial positions (initialization = “random”). The diffusion coefficients of particles both inside and outside were set to 1 µm^2^/s (diffConstIn = 1; diffConstOut = 1). The time step was set to 0.03 s (dt = 0.03). The domain and nuclear boundaries were reflective according to the classical Snell-Descartes scheme for reflection (boundary = “reflective”). We ran 200 realizations as ground-truth datasets and analyzed them using “AnalyzeTrajConvexHull” to compare the classification. Note that “AnalyzeTrajConvexHull” exports results of classification based on various convex hull statistics, including perimeter, area, and maximum diameter. Here we compared the results of perimeter with the ground-truth and calculated the classification accuracy and the confusion matrix. To determine how the domain radius to be detected and the particle diffusion coefficient affect the accuracy of the fitted radius, we additionally simulated scenarios of varying domain radii (from 0.1 µm to 2.5 µm) and diffusion coefficients (from 0.5 µm^2^/s to 2 µm^2^/s). Then, we computed the accuracy (1 - percentage error) of the fitted radius compared to the simulated domain radius. Specific details of the algorithms used are described below:

#### Step 1: Simulation of Brownian trajectories

We generated 2D trajectories with positions *X*(*t*) ∈ R^2^ at time *t*. Each trajectory has *K* + 1 points with a fixed time step Δ*t*. The result is the points *X*(0),*X*(Δ*t*),…,*X*(*K*Δ*t*). By using Euler’s scheme of the Smoluchowski’s limit equation (Schuss 2010; Holcman and Schuss 2017), the Brownian motion of the trajectories was given by:

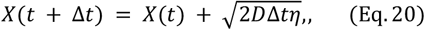

where *D* is the diffusion coefficient, *η* is a Gaussian variable with mean 0 and variance 1.

#### Step 2: Modelling trajectories inside a Phase Separated Domain (PSD)

To model trajectories confined by a PSD, we constructed two concentric discs, where the inner disc Ω_PSD_ represented the nuclear PSD, and the outer disc Ω_nuc_ the nucleus. The radius of the PSD, *R*_PSD_, was smaller than that of the nucleus, *R*_nuc_. To ensure that trajectories inside the PSD cannot escape and trajectories outside cannot enter, we made the boundaries of the two domains *∂*Ω_PSD_ and *∂*Ω_nuc_ reflective on both sides. We then ran simulations after setting the number *m*_PSD_ and *m*_nuc_ of trajectories inside and outside the PSD, respectively. The initial positions *X*(0) were uniformly distributed in the two domains. We then generated Brownian trajectories from (Eq.20). At each step, if a trajectory crossed the boundaries, it was reflected using the classical Snell-Descartes scheme against the normal to the boundaries. We computed the convex hull perimeter from the simulated trajectories as described above and used them as ground-truth, where the trajectories inside the PSD were labelled as domain-confined (𝒞) and the outside trajectories as unconfined (𝒰). The ground-truth label *T*(*i*) of trajectory *i* was thus defined by:

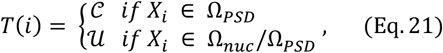

#### Step 3: Evaluating the classifier using the ground-truth trajectories

To measure the accuracy of our computational pipeline, we classified the ensemble of simulated domain-confined and unconfined trajectories, which assigned each simulated trajectory *i* with a label *P*(*i*) ∈ {𝒞, 𝒰}. For one realization, we computed the accuracy *M* of the classification compared to the ground-truth labels *T*(*i*) by:

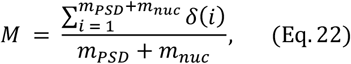

where *δ*(*i*) is an indicator function defined by:

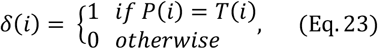

We averaged the classification accuracy over *N* realizations: 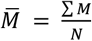. The confusion matrix was computed by averaging the number of true-positives (*P* = *T* = 𝒞), true-negatives (*P* = *T* = 𝒰), false-positives (*P* = 𝒞, *T* = 𝒰) and false-negatives (*P* = 𝒰, *F* = 𝒞) over the *N* realizations. Finally, we also estimated the radius 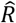 fit and its accuracy:

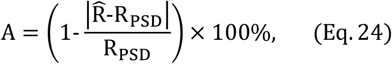

where we used the chosen radius *R*_PSD_ and the fitted one 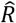. The average was taken over *N* realizations: 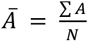.

### Simulations using empirical estimates from the TF datasets

To understand whether constriction by a domain reduces anomalous exponent, we used the “AggregateDiffusionSimulator” to simulate domain-confined and unconfined trajectories and calculated their anomalous exponents. Simulation parameters were set according to the empirical estimates of the TFs:

| Parameter | Parameter description | NANOG-WT | SOX2 | NANOG-W10A | Unit |
| --- | --- | --- | --- | --- | --- |
| dt | Time step | 0.03 | 0.03 | 0.03 | s |
| $R_{\text{PSD}}$ | Domain radius | 0.39 | 0.445 | 0.325 | $\mu\text{m}$ |
| $R_{\text{nuc}}$ | Nuclear radius | 10 | 10 | 10 | $\mu\text{m}$ |
| $m_{\text{PSD}}$ | #Particles inside domain | 610 | 640 | 660 | particles |
| $m_{\text{nuc}}$ | #Particles outside domain | 390 | 360 | 340 | particles |
| D | Diffusion coefficient | 0.91 | 0.99 | 0.97 | $\mu\text{m}^2/\text{s}$ |
| nStep (K) | Trajectory length | 5 | 5 | 5 | steps |

## Author contributions

**Conceptualization:** SBa, TEB, IOdeA. **Data curation**: DJ, YC. **Formal Analysis:** DJ, YC with help from SES, GGA. **Funding acquisition:** SBa, DH, SFL, TEB, SES, AW, KJC. **Investigation:** GGA, DJ, SD. **Methodology:** YC, SES, GGA, DJ, SD with help from SM. **Project administration:** GGA, SBa, DH. **Resources:** (cell lines) AW, GGA, MJ, BDH with help from LB. (SMLM instrument building/maintenance) SD, SFL, ZZ, SBa, DK. **Software:** YC, SES, DJ with help from GN, SBe, KB. **Supervision:** SBa, DH, TEB, RP-C, KJC. **Validation:** (Cell lines) GGA, KG, AW, SES, TP, JAN. **Visualization:** DJ, YC, SES. **Writing:** SBa, DH with help from DJ, YC, GGA. **Writing – review & editing:** SBa, SES, RP-C.

## Acknowledgements

We thank Jenny Nichols and all members of the Basu and Holcman labs for insightful and critical discussions. We thank Robin Floyd for help making the NANOG-HALO ESCs and Tomas Hanak for assembling the Photometrics Python 3D fitting code. We thank Darran Clements in the Cambridge Stem Cell Institute (CSCI) Imaging Facility, and David Gaboriau and Volodymyr Nechyporuk-Zloy in the Imperial College London (ICL) Facility for Imaging by Light Microscopy (FILM) for training students/staff and assisting with experiments. We thank Fiona Watt, Sally Lees and the CSCI Tissue Culture Core Facility, and Javaid Iqbal in the ICL Department of Life Sciences (DoLS) Tissue Culture Core Facility for technical advice and support.

GGA was funded by the UKRI Biotechnology and Biological Sciences Research Council (BBSRC) (BB/W000423/1, BB/W000423/2). YC and DH are supported by grants from the ANR (ANR-24-CE12-5087 - InsulatorGrammar, ANR-22-CE16-0027 AstroXcite) and the European Research Council (ERC) under the European Union’s Horizon 2020 research and innovation programme (grant agreement No 882673). SES was funded by the Joint Research Grant (Isaac Newton Trust, Wellcome Trust ISSF, University of Cambridge), the CSCI Wellcome Trust seed fund, the Wellcome Trust Sir Henry Wellcome Fellowship (224070/Z/21/Z) and a University of Sheffield Strategic Research Fellowship in the Physics of Life and Quantitative Biology. SD was funded by the BBSRC (BB/X511092/1 and UKRI715). ZZ was funded by the UKRI Engineering and Physical Sciences Research Council (EPSRC) (EP/W015005/1). AW was funded by the Wellcome Trust PhD programme in Stem Cell Biology and Medicine (218481/Z/19/Z). LB was funded by the UKRI Medical Research Council (MRC) (MR/R017735/1). TP and JAN were funded by the Wellcome Trust (215477/Z/19/Z). SBa was part-funded by the Trinity College Stem-Cell Medicine Senior Postdoctoral Researcher Fund and a starting grant at the Cambridge Stem Cell Institute from the Wellcome Trust (203151/Z/16/Z) and UKRI MRC (MC_PC_17230). RP-C and SM were funded by the Leverhulme Trust Research Project (RPG-2023-085). RP-C was also funded by the BBSRC (BB/Y002709/1). The Facility for Imaging by Light Microscopy (FILM) at Imperial College London is part-supported by funding from the Wellcome Trust (104931/Z/14/Z) and BBSRC (BB/L015129/1 and BB/T017929/1).

